# Mitochondria branch within Alphaproteobacteria

**DOI:** 10.1101/715870

**Authors:** Lu Fan, Dingfeng Wu, Vadim Goremykin, Jing Xiao, Yanbing Xu, Sriram Garg, Chuanlun Zhang, William F. Martin, Ruixin Zhu

**Affiliations:** Academy for Advanced Interdisciplinary Studies, Southern University of Science and Technology (SUSTech), Shenzhen 518055, China; Shenzhen Key Laboratory of Marine Archaea Geo-Omics, Department of Ocean Science and Engineering, Southern University of Science and Technology (SUSTech), Shenzhen 518055, China; Putuo people’s Hospital, School of Life Sciences and Technology, Tongji University, Shanghai 200092, P.R.China; Research and Innovation Centre, Fondazione E. Mach, 38010 San Michele all’Adige (TN), Italy; Laboratory for Marine Geology, Qingdao Pilot National Laboratory for Marine Science and Technology, Qingdao, 266061, China; Institute of Molecular Evolution, Heinrich-Heine-University, Universitätsstr. 1, 40225 Düsseldorf, Germany

## Abstract

It is well accepted that mitochondria originated from an alphaproteobacterial-like ancestor. However, the phylogenetic relationship of the mitochondrial endosymbiont to extant alphaproteobacteria remains a subject of discussion. The focus of much debate is whether the affiliation between mitochondria and fast-evolving alphaproteobacterial lineages reflects true homology or artifacts. Approaches such as protein-recoding and site-exclusion have been claimed to mitigate compositional heterogeneity between taxa but this comes at the cost of information loss and the reliability of such methods is so far unjustified. Here we demonstrate that site-exclusion methods produce erratic phylogenetic estimates of mitochondrial origin. We applied alternative strategies to reduce phylogenetic noise by taxon replacement and selective exclusion while keeping site substitution information intact. Cross-validation based on a series of trees placed mitochondria robustly within Alphaproteobacteria.

## Introduction

The origin of mitochondria is one of the defining events in the history of life. Although alternative explanations do exist (e.g. the mosaic origin ^1^), gene-network analyses ^2–5^ and marker gene-based phylogenomic inference (see review by Roger et al. ^6^) have generally reached a consensus that mitochondria have a common bacterial ancestor, which was a close relative to extant alphaproteobacteria. However, the exact relationship of mitochondria to specific alphaproteobacterial groups remains contentious. Phylogenetic placement of mitochondria in the tree of Alphaproteobacteria has been extremely difficult for several reasons.

They include considerable phylogenetic divergence and metabolic variety within Alphaproteobacteria ^2-5,7^, faint historical signals left behind the very ancient event of mitochondria origin ^8^, limited number of marker genes shared between mitochondria and Alphaproteobacteria due to extensive gene loss in the prior ^9^, taxonomic bias in datasets towards clinically or agriculturally important alphaproteobacterial members ^10^. Furthermore, these effects are compounded by strong phylogenetic artifacts associating mitochondria with some fast-evolving alphaproteobacterial lineages such as Rickettsiales and Pelagibacterales resulting in erroneous clade formations (see a detailed review in Roger et al. (2017)).

To minimize the possible influence of long-branch attraction coupled with convergent compositional signals, various strategies have been applied such as the use of nucleus-encoded mitochondrial genes ^5,11,12^, site or gene exclusion ^13–15^, protein recoding ^15^ and the use of heterogeneity-tolerant models such as the CAT model implemented in Bayesian inference ^11,16^. These attempts have generally proposed four hypotheses: (1) mitochondria root in or as the sister of Rickettsiales ^12,17^), which are all obligate endosymbionts (but see reference ^18^); (2) mitochondria are sisters with free-living alphaproteobacteria such as *Rhodospirillum rubrum* ^14^, Rhizobiales and Rhodobacterales ^5^; (3) mitochondria are neighbors to a group of uncultured marine bacteria ^10^; and (4) mitochondria are most closely related to the most abundant marine surface alphaproteobacteria – SAR11 (referred as Pelagibacterales in this study) ^19,20^. While the first hypothesis has been reported most frequently so far, the last has been explained by several independent groups as a result of compositional convergence artifact ^10,13,16^.

Recently, Martijn et al. revisited this topic by using a dataset including alphaproteobacterial genomes assembled from the Tara Ocean metagenomes ^21^. They reported that when compositional heterogeneity of the protein sequence alignments was sufficiently reduced by site exclusion and to fit their specified model, the entire alphaproteobacterial class formed a sister group to mitochondria. Their conclusion challenged the long-agreed phylogenetic consensus that mitochondria originated from within the Alphaproteobacteria ^22^. However, model over-fitting comes at a cost of information loss and does not guarantee correct phylogenetic prediction. While excluding possible noise in compositionally heterogenous sites might mitigate systematic errors, it can also lead to model overfitting. *A priori*, one cannot rule out the possibility that these sites contain phylogenetic information of true evolutionary connection between mitochondria and Alphaproteobacteria? A similar concern about information loss and a demand for further justification of their results was also voiced by Gawryluk ^23^.

We here examined the phylogenetic affiliations of mitochondria by using several site-exclusion methods and demonstrated that these results should be interpreted with utmost caution. We then applied a different approach to significantly reduce compositional signals in the dataset by taxon replacement and selectively lineage exclusion while keeping the native site substitution intact. We successfully resolved relationship of fast-evolving lineages including mitochondria with slowly-evolving alphaproteobacteria. Our results support the traditional view that mitochondria branch within Alphaproteobacteria.

## Results

### Site exclusion approaches produced stochastic phylogenetic inference for mitochondria

The idea of excluding potentially model-violating sites to improve phylogenetic prediction was introduced over two decades ago ^24,25^ but has been opposed by researchers (see review by Shepherd et al. ^26^). The concern is that in spite of non-historical signals, these sites may contain useful information. Nonetheless, various versions of site exclusion have been applied in phylogenetic studies of mitochondria and Alphaproteobacteria either based on evolving rate ^14,15^ or amino acid composition ^13,21,27^. However, conflicting results were reported by using different site-exclusion metrics ^15^.

To cross-validate the effects of site-exclusion approaches on mitochondrial and alphaproteobacterial phylogeny, we implemented five metrics with different principles in this study (Table 1). Among them, Stuart’s test and Bowker’s test are two typical evaluation metrics of symmetry violation ^28^. Compared to Stuart’s test, Bowker’s test of symmetry was reported to more comprehensive and sufficient to assess the compliance of symmetry, reversibility and homogeneity in time-reversible model assumptions ^28^. The χ^2^-score metric was designed to test site contribution to dataset compositional heterogeneity ^13^ and was applied by Martijn et al. for mitochondrial phylogeny study ^21^. ɀ-score is a metric specifically designed to cope with strong GC content-related amino acid compositional heterogeneity in datasets of alphaproteobacterial phylogeny ^27^. A method implemented in IQTREE for fast-evolving site selection was also included for comparison since long-branch attraction caused by fast-evolving species in Alphaproteobacteria and mitochondria is a potential issue ^29^.

**Table 1.**
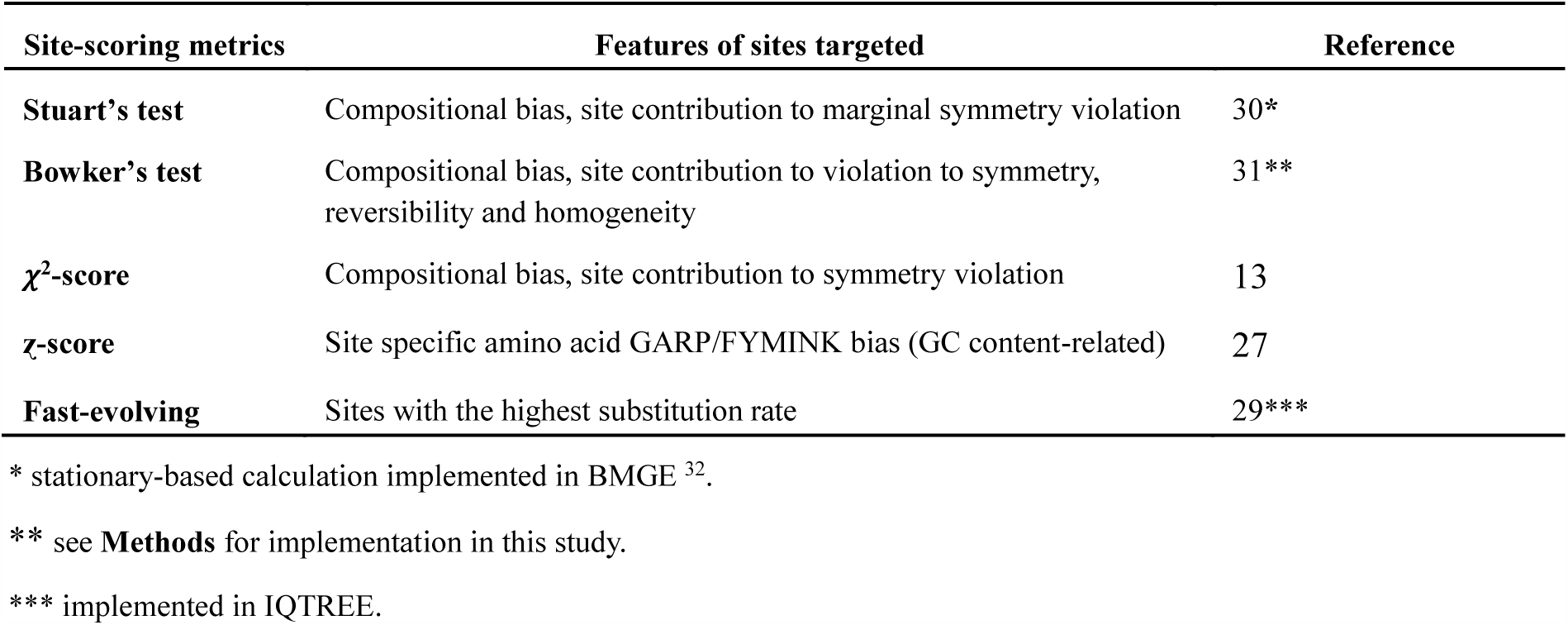
Introduction of site-exclusion methods for justification.

| Site-scoring metrics | Features of sites targeted | Reference |
| --- | --- | --- |
| **Stuart's test** | Compositional bias, site contribution to marginal symmetry violation | 30* |
| **Bowker's test** | Compositional bias, site contribution to violation to symmetry, reversibility and homogeneity | 31** |
| **$\chi^2$-score** | Compositional bias, site contribution to symmetry violation | 13 |
| **ζ-score** | Site specific amino acid GARP/FYMINK bias (GC content-related) | 27 |
| **Fast-evolving** | Sites with the highest substitution rate | 29*** |

\* stationary-based calculation implemented in BMGE<sup>32</sup>.
\*\* see **Methods** for implementation in this study.
\*\*\* implemented in IQTREE.

Site-excluded subsets of the ‘24-alphamitoCOGs’ dataset in Martijn et al. (2018) were generated by using the five methods with a series of cutoff values except for Stuart’s test on which a single stationary-base calculation was applied (**Supplementary Table 1**). Trees of the subsets were compared to the tree of the untreated dataset, respectively. Topological dissimilarity between two trees was calculated by using the Alignment metric ^33^. This method was found to superior among other tree comparison metrics ^34^. Both simple model and mixed model (C60) were used in Maximum-likelihood (ML) tree reconstruction for comparison (tree files are deposited in **Supplementary Data Files**). Site exclusion approaches led to substantial tree topological changes (Fig. 1). In general, the increase in number of sites removed precipitated increases in changes of tree topology. Among the five methods, ɀ-score generally caused the least changes in nearly all the subsets of alignment. These patterns are consistent when either simple or mixed models were applied in phylogenetic inference.

**Fig. 1.**
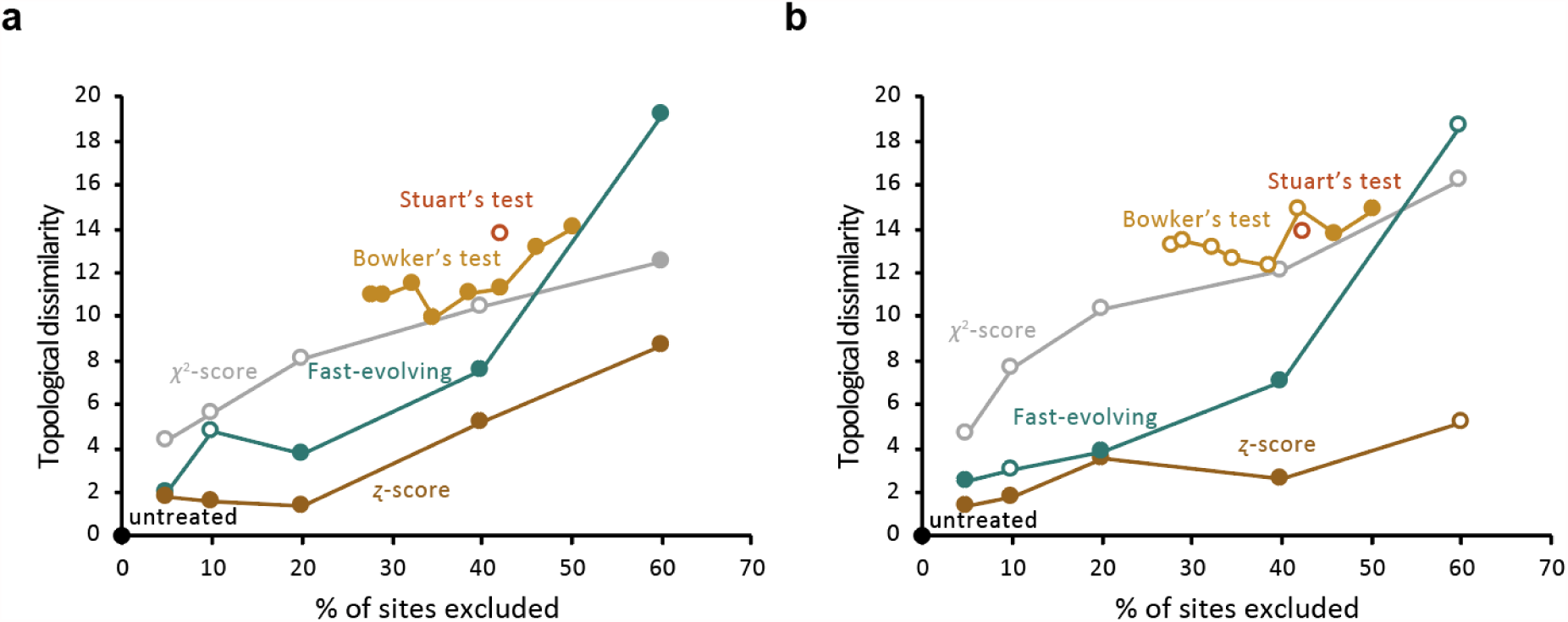
Tree dissimilarity based on the Alignment metric between the untreated tree and trees generated after applying site-exclusion approaches. All trees are rooted. Empty dots show trees supporting the Alphaproteobacteria-sister topology and filled dots show trees supporting the within-Alphaproteobacteria topology of mitochondria. **a**, ML trees under simple models. **b**, ML trees under the mixed model (C60).

We summarized the position of mitochondria in these site-excluded trees and stochastic results were observed (Fig. 1). Nearly half of the trees support mitochondria in a sisterhood with the entire Alphaproteobacteria (‘mito-out’) and the other half support that mitochondria branch within Alphaproteobacteria (‘mito-in’). Noticeably, while we reproduced the results observed in Martijn et al. (2018) that tree topology shifted from ‘mito-in’ to ‘mito-out’ when 5% to 40% of sites were removed by using the *χ*^2^-score metric, exclusion of more sites (60% here) change the tree topology back to ‘mito-in’ predicted by the simple model (Fig. 1a). It is likely that site-exclusion method, the number of sites excluded and tree model applied had a mixed function to the phylogenetic relationship of mitochondria to Alphaproteobacteria. One explanation to this observation is that sites strongly supporting either the ‘mito-in’ or the ‘mito-out’ topology were randomly excluded by these metrics. The absence of certain topology-determining and ‘mito-in’-supporting sites can cause tree shift from one topology to the other, while the further loss of ‘mito-out’-supporting sites may shift the tree topology back.

### Taxa replacement efficiently reduced compositional heterogeneity between lineages of interest

To counter compositional heterogeneity but without arbitrarily compromising phylogenetic signals, we then replaced the mitochondrial and Rickettsiales sequences with GC-rich alternatives. Specifically, while keeping most of the taxa used in the ‘24-alphamitoCOGs’ dataset (see **Methods**), five less AT-rich mitochondria (GC content 45.1%-52.2% compared to 22.3%-40.6% in the original dataset) and five less AT-rich Rickettsiales (GC content 38.2%-49.8% compared to 29.0%-50.0% in the original dataset) were selected to replace the mitochondrial and rickettsiales groups in the original dataset (**Supplementary Table 2**). The GC-poor vs. GC-rich amino acid (FYMINK/GARP) ratio of marker proteins of the reselected mitochondria and Rickettsiales ranged from 0.955 to 1.329 and from 1.013 to 2.330, respectively (Fig. 2). In comparison to the ‘24-alphamitoCOGs’ dataset, we have remarkably reduced the heterogeneity in FYMINK/GARP ratio between mitochondria and slowly-evolving alphaproteobacteria.

**Fig. 2.**
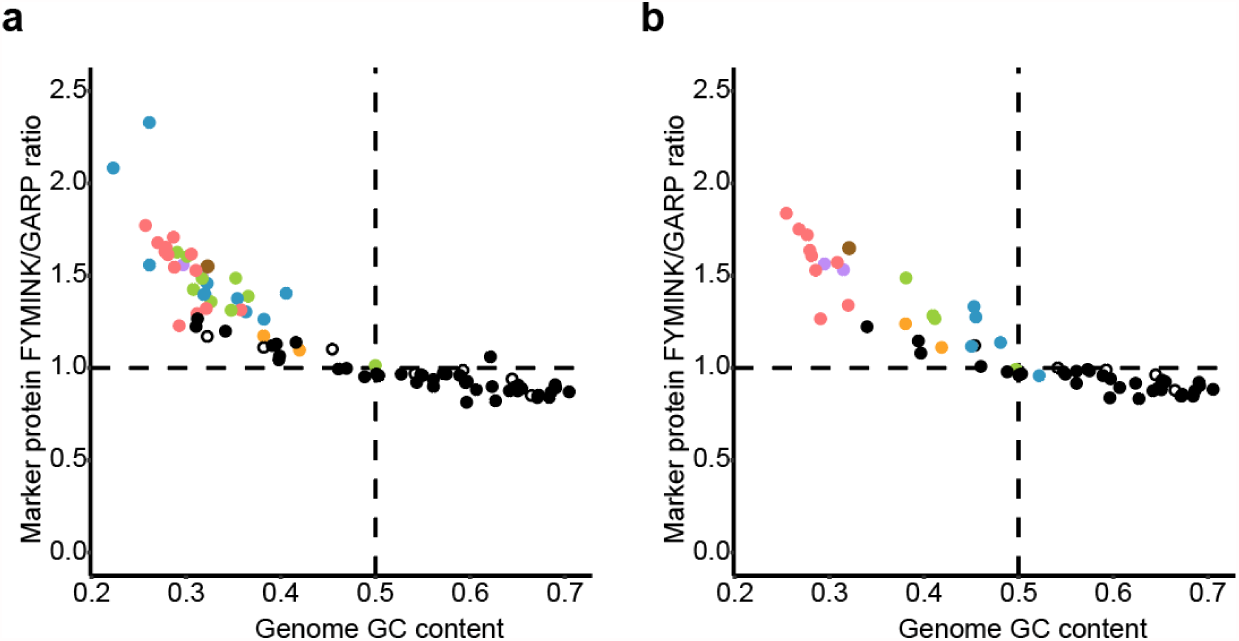
GC content and amino acid compositional heterogeneity among alphaproteobacterial lineages and mitochondria. Dots represent taxa. Lineages are colored according to Fig. 4 except empty dots represent Beta-, Gammaproteobacteria and Magnetococcales. **a**, taxa in the ‘24-alphamitoCOGs’ dataset. **b**, taxa in the ‘18-alphamitoCOGs’ dataset.

In total, 61 nonredundant taxa were selected and 18 of the original 24 marker proteins were used for phylogenetic inference (**Supplementary Table 2, 3**). We named our new dataset ‘18-alphamitoCOGs’. It is needed to notice that the introduced GC-rich mitochondria were all from higher plants. While this may compromise the representation of data, the mitochondrial sequences of higher plants are considered to have diverged from bacterial sequences to the least extent ^35,36^ as a result of low mutation rate in genes possibly maintained by DNA repair mechanisms ^37^.

### Taxon-reduced datasets produced congruent phylogenetic prediction for fast-evolving alphaproteobacteria

A meaningful alphaproteobacterial species phylogeny is prerequired in investigating the phylogenetic relationship between Alphaproteobacteria and mitochondria. However, until recently, the tree topology of Alphaproteobacteria is not yet fully resolved ^27^. We tested our new dataset for resolving phylogeny between alphaproteobacterial lineages. First, to minimize the interference between fast-evolving lineages in the same tree, fast-evolving taxa were excluded for ML and Bayesian tree reconstruction (see **Methods**). The remaining alphaproteobacteria are expected to contain minimum non-historical signals and less likely cause model violation. While the ML tree and Bayesian tree were slightly different in the topology of basal branches, they reached an agreement that these slowly-evolving alphaproteobacteria can be classified into four major clades, which were named as Alpha I, Alpha II, Alpha III and GT, respectively (Fig. 3ab, **Supplementary Fig. 1, 2**, **Supplementary Table 2**). We here assign these alphaproteobacteria as ‘backbone taxa’ and the four clades as ‘backbone clades’.

**Fig. 3.**
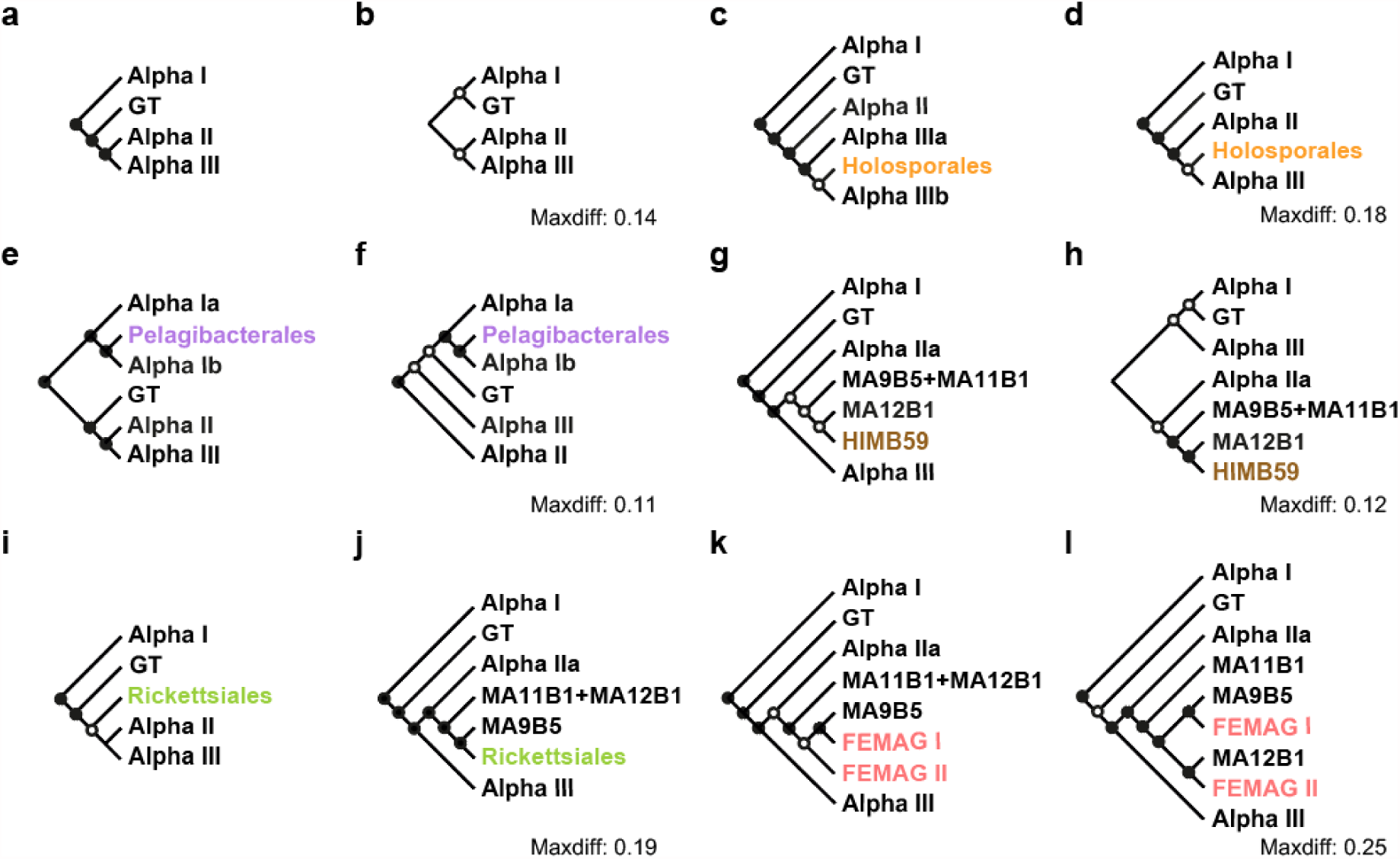
Schematic phylogenetic trees of subgroups of Alphaproteobacteria in the ‘18-alphamitoCOGs’ dataset. Alphaproteobacterial lineages are named according to **Supplementary Table 2**. Taxa and taxonomic groups in black present the backbone taxa. Filled dots show node support values greater than 80% while empty dots show values greater than 50% but less than 80%. Node values show posterior probability support values for Bayesian trees and bootstrapping support values based on 1000 iterations for ML trees. Trees are rooted. Outgroup taxa and *Magnetococcus marinus* MC-1 are not shown. The Maxdiff values of Bayesian trees are shown beside the trees. **a-l**, Schematic trees of **Supplementary Fig. 1-12**, respectively.

Group GT is equivalent to Geminicoccaceae in Muñoz-Gómez et al. (2019) ^27^. Alpha I comprises core Alphaproteobacterial orders including Kordiimonadales, Sphingomonadales, Rhizobiales, Caulobacterales, Parvularculales and Rhodobacterales. Grouping of these lineages is in consistence with the findings by Muñoz-Gómez et al. and others ^7,27,38^. Alpha II comprises three isolates belonging to Rhodospirillaceae and several marine alphaproteobacterial metagenome-assembled genomes (MAGs). Grouping of these lineages was observed in by Williams et al. and others ^7,27,38^. Alpha III comprises Kiloniellaceae, SAR116, Acetobacteraceae, Azospirillaceae, and some taxa classified to the polyphyletic Rhodospirillaceae. This result is similar to the finding by Muñoz-Gómez et al. ^27^. Noticeably, separation of these four groups were exactly recovered by Martijn et al. in their untreated ‘24-alphamitoCOGs’ dataset (Supplementary Fig. 9, 10 in Martijn et al. (2018)), but not in their stationary-trimmed dataset (Fig. 4a and Supplementary Fig. 11, 12 in Martijn et al. (2018)), which they claimed to support their ‘mito-out’ result. This again suggests site-exclusion may result in abnormal tree topology for even slow-evolving species.

**Fig. 4.**
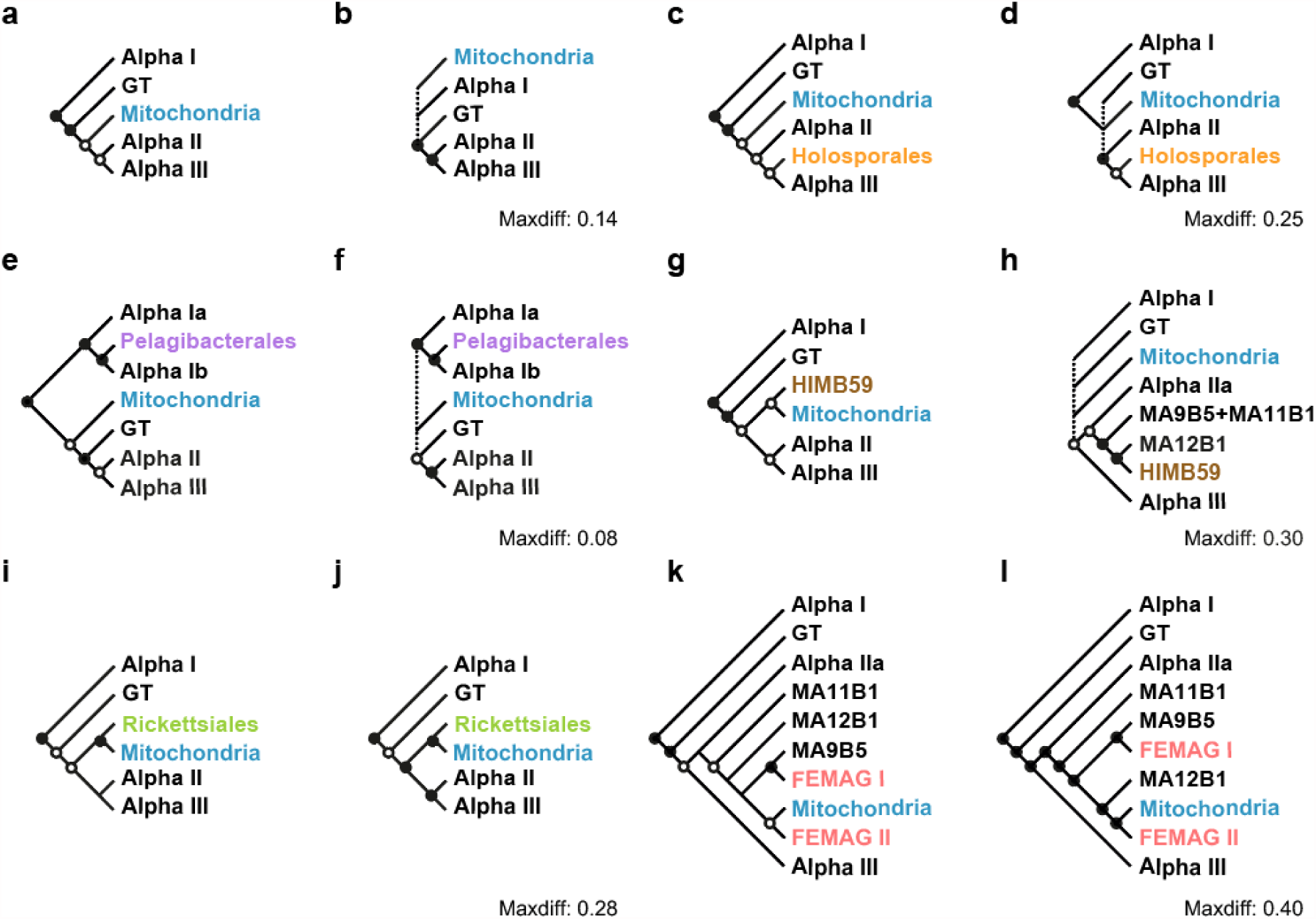
Schematic phylogenetic trees of mitochondria and subgroups of Alphaproteobacteria in the ‘18-alphamitoCOGs’ dataset. Alphaproteobacterial lineages and Mitochondria are named according to **Supplementary Table 2**. Taxa and taxonomic groups in black present the backbone taxa. Filled dots show node support values greater than 80% while empty dots show values greater than 50% but less than 80%. Node values show posterior probability support values for Bayesian trees and bootstrapping support values based on 1000 iterations for ML trees. Trees are rooted. Outgroup taxa and *Magnetococcus marinus* MC-1 are not shown. The Maxdiff values of Bayesian trees are shown beside the trees. **a-l**, Schematic trees of **Supplementary Fig. 13-24**, respectively.

Each of the six groups of fast-evolving alphaproteobacteria were then added to this dataset of backbone taxa and a series of phylogenetic trees were built, respectively. Despite the addition and removal of fast-evolving taxa, backbone taxa maintained a topology in which all the four backbone clades maintained their monophyly (Fig. 3).

Holosporales was previously considered as a subclade of Rickettsiales based on phylogenetic analysis and the factor that members of both groups are obligate endosymbionts (but see **Discussion**) ^39–41^. However, other studies suggested the phylogenetic affiliation between Holosporales and other Rickettsiales families are the result of artifact ^12,20,21,39,42^. In the very recent study by Muñoz-Gómez et al. where amino acid bias was corrected by site exclusion, Holosporales was suggested to have a derived position within the Rhodospirillales and possibly close to Azospirllaceae ^27^. In our study, the ML tree suggests Holosporales be placed in Alpha III forming a sister relationship with Azospirllaceae and Acetobacteraceae (Alpha IIIb) with a weak node support, while the Bayesian result suggests they are in sisterhood with the entire Alpha III (Fig. 3cd, **Supplementary Fig. 3, 4**). Our results suggest that Holosporales are distant to Rickettsiales but close to taxa in Alpha III.

Recent studies suggested the grouping of Pelagibacteriales, alphaproteobacterium HIMB59 and Rickettsiales, as reported by many earlier studies is the result of a compositional bias artefact ^21,27^. Using site-exclusion datasets, it was suggested that Pelagibacteriales should be placed after the common ancestor of Sphingomonadales (belonging to Alpha Ia here) but before the divergence of Rhodobacterales, Caulobacterales and Rhizobiales (belonging to Alpha Ib here) ^27^. Our result is consistent with this (Fig. 3ef, **Supplementary Fig. 5, 6**). Moreover, without interference of other fast-evolving species, alphaproteobacterium HIMB59 here was placed in the clade Alpha IIb forming a sisterhood with MarineAlpha 12 Bin1 (Fig. 3gh, **Supplementary Fig. 7, 8**).

**Fig. 5.**
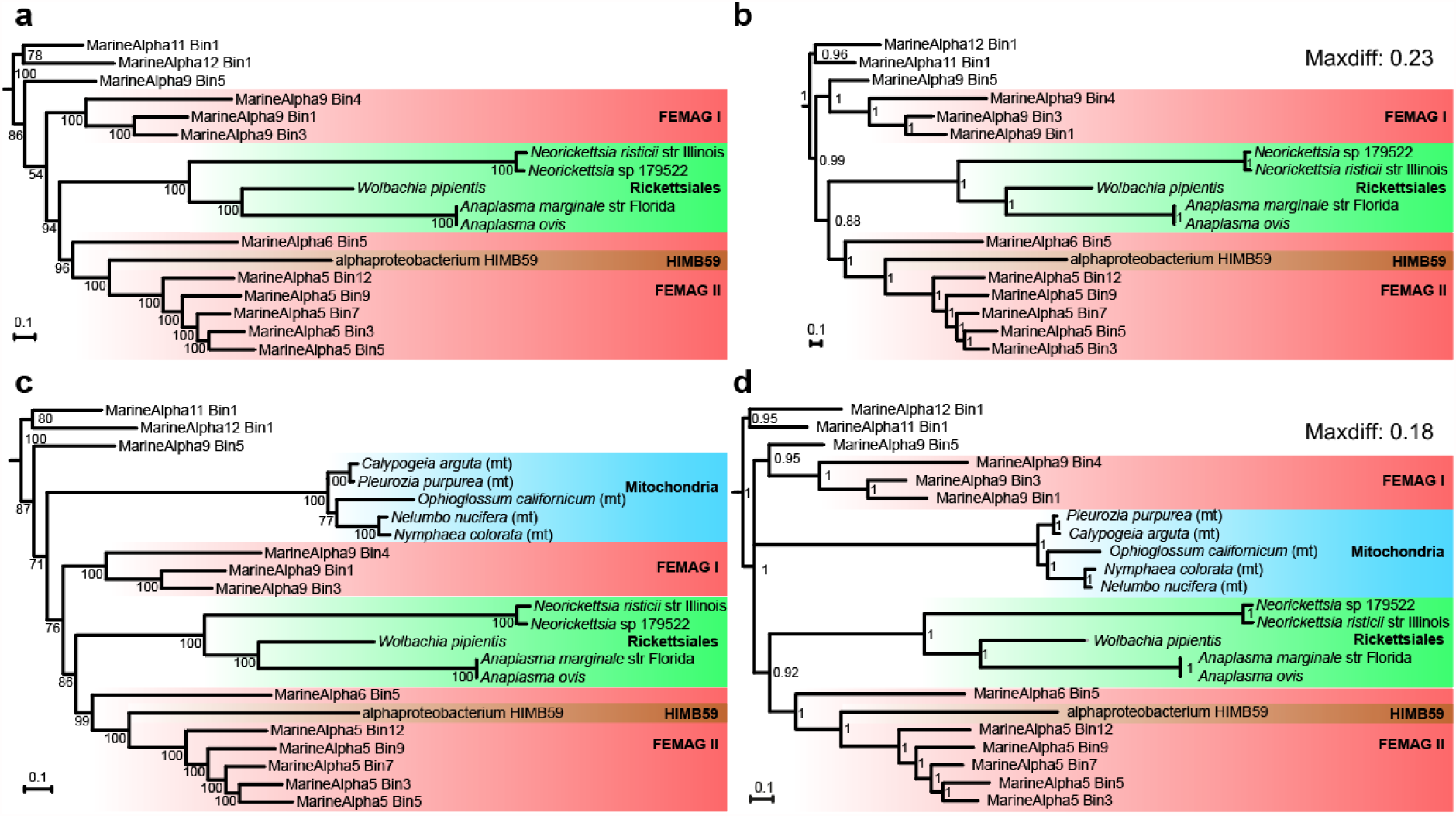
Phylogenetic relationships of fast-evolving taxa and mitochondria to alphaproteobacteria of Alpha IIb. Node values show posterior probability support values for Bayesian trees and bootstrapping support values based on 1000 iterations for ML trees. mt, mitochondria. All trees are rooted and the outgroup is not shown. **a** and **b**, ML and Bayesian trees, respectively, of fast-evolving alphaproteobacteria and taxa of Alpha IIb. **c** and **d**, ML and Bayesian trees, respectively, of fast-evolving alphaproteobacteria, mitochondria and taxa of Alpha IIb.

Rickettsiales appearing as sister to all other alphaproteobacteria has been reported in some artifact-attenuated studies ^27^ while conflicting results were recovered in others ^21^ suggesting the current difficulty in resolving its relationship with slow-evolving alphaproteobacteria. We found that Rickettsiales were placed as sister to the clade of Alpha II and Alpha III in the ML tree with a weak basal node support (Fig. 3i, **Supplementary Fig. 9**). Interestingly, however, in the converged Bayesian tree, Rickettsiales was placed within Alpha II, as the sister of MarineAlpha9 Bin5, suggesting possible connection between Rickettsiales and this newly discovered, non-fast-evolving marine alphaproteobacterium (Fig. 3j, **Supplementary Fig. 10**).

Including MarineAlpha9 Bin5, Martijn et al. obtained a number of marine alphaproteobacterial MAGs from the Tara Oceans project ^43^. In both ML and Bayesian trees, fast-evolving MAGs belonging to FEMAG I and FEMAG II were robustly placed within Alpha IIb (Fig. 3kl, **Supplementary Fig. 11, 12**). Specifically, FEMAG I showed a strong connection to MarineAlpha9 Bin5, while FEMAG II was linked to MarineAlpha12 Bin1 in the Bayesian tree.

### Taxon replacement and selective exclusion approaches placed mitochondria within Alphaproteobacteria

To study the phylogenetic relationship of mitochondria to alphaproteobacterial groups, we added GC-neutral mitochondria to the trees of backbone taxa solely or in combinations with other fast-evolving clades. Mitochondria by themselves were placed within Alphaproteobacteria as the sister of Alpha II and Alpha II in the ML tree with a weak node support (Fig. 4a, **Supplementary Fig. 13**). However, the counterpart Bayesian tree could not resolve the relationship of mitochondria to taxa of the four alphaproteobacterial backbone clades (Fig. 4b, **Supplementary Fig. 14**). Similar results were observed in trees including mitochondria in combination with Holosporales, Pelagibacterales and alphaproteobacterium HIMB59, respectively (Fig. 4c-h, **Supplementary Fig. 15-20**). Specifically, in ML trees, mitochondria were always placed within Alphaproteobacteria with low bootstrap support (71%, 57%, and 65%, respectively). In Bayesian trees, none of the three fast-evolving clades could provide adequate information in resolving the phylogeny of mitochondria. Our approach successfully broke the frequently reported false grouping of Holosporales, Pelagibacterales, alphaproteobacterium HIMB59 and mitochondria causing by compositional convergence and clearly suggested that there is little phylogenetic connection between mitochondria and these three alphaproteobacterial lineages.

In contrast, apparent phylogenetic connection of mitochondria to Rickettsiales and FEMAG II were observed in both ML and Bayesian trees (Fig. 4i-l, **Supplementary Fig. 21-24**). Specifically, mitochondria and Rickettsiales were placed together independently to the four backbone clades (node support 97% for the ML tree), while mitochondria and FEMAG II were placed in sisterhood inside the Alpha IIb clade (node support 68% for the ML tree). The inconsistence in the relative placement of mitochondria to the backbone clades could be the result of insufficient taxon sampling.

### Phylogenetic relationships of taxa in clade Alpha IIb provided novel insights into the origin of mitochondria

Since Rickettsiales, alphaproteobacterium HIMB59, FEMAG I and FEMAG II individually showed phylogenetic connections to taxa of Alpha IIb in Bayesian trees, evolutionary relationships between these lineages were then investigated specifically by setting Alpha IIa (**Supplementary Table 2**) as the outgroup. MarineAlpha11 Bin1 and MarineAlpha12 Bin2 formed a monophylic clade in both trees (Fig. 5ab). MarineAlpha9 Bin5 either branched below all the fast-evolving taxa studied here in the ML tree or formed monophyly with FEMAG I in the Bayesian tree. The nodes connecting the branch of MarineAlpha9 Bin5 and the branch of FEMAG I, respectively, had low support suggesting the phylogenetic relationship between these two branches in the ML tree was unstable. Both trees reached an agreement that alphaproteobacterium HIMB59 branched within FEMAG II and Rickettsiales was in sisterhood with FEMAG II. This result suggests both Rickettsiales and alphaproteobacterium HIMB59 are evolutionarily connected to a group of uncultured marine planktonic alphaproteobacteria.

Moreover, as mitochondria showed strong phylogenetic connections to both Rickettsiales and FEMAG II (Fig. 4), we then included mitochondria in these two trees. When mitochondria were present, the topology of all other taxa was preserved in both the ML tree and the Bayesian tree (Fig. 5cd). Mitochondria were placed below the clade consist of FEMAG II, alphaproteobacterium HIMB59, Rickettsiales and FEMAG I in the ML tree with node support of 71%. In comparison, the phylogenetic relationship of mitochondria, the clade of FEMAG I and MarineAlpha9 Bin5 and the clade of FEMAG II, alphaproteobacterium HIMB59 and Rickettsiales was unresolved by Bayesian inference. Despite that, the placement of mitochondria within Alpha IIb was robust. Our result suggests that mitochondria may have originated from the common ancestor of Rickettsiales and certain extant marine planktonic alphaproteobacteria.

The placement of mitochondria together with fast-evolving taxa within Alpha IIb is unlikely a result of phylogenetic artifact based on several lines of evidence. First, taxon-exclusion analyses clearly demonstrate the phylogenetic connections of these fast-evolving alphaproteobacterial lineages to non-fast-evolving taxa MarineAlpha9 Bin5 and MarineAlpha11 Bin1 in the absence of possible influence from non-historical signals (Fig. 3). Secondly, in our analysis, mitochondria and these fast-evolving taxa did not form a singlet clade falling apart from backbone clades as a result of long-branch attraction – something shown in Supplementary Fig. 9, 10 in Martijn et al. (2018). Instead, they were placed together with slowly-evolving taxa within Alpha IIb. Lastly, there were divergent FYMINK/GARP ratios among Rickettsiales, mitochondria and FESMASs (Fig. 2). A compositional convergence artifact would actually have separated them instead of grouped them.

## Discussion

As datasets in studies on phylogeny between mitochondria and Alphaproteobacteria heavily suffer from compositional heterogeneity and long-branch attraction, various approaches to mitigate non-historical signals have been adopted but the drawbacks of these methods are rarely examined. Among them, protein recoding cause signal loss and artificial mutation saturation ^44^. Nucleus-encoded mitochondrial genes have to be adapted to new rules of expression and regulation in the nucleus system and therefore may actually have undergone intensive site substation compared to mitochondrion-encoded genes. Thus, the reliability of using nucleus-encoded mitochondrial genes in phylogenetic analysis of mitochondria need further justification ^11,45^. In this study, we further demonstrated that site-exclusion methods can impair the study of mitochondrial phylogeny by causing random topological shifts, particularly among basal branches, via arbitrary cutoff selection, thereby breaking well-established phylogenetic relationships of even homogeneous datasets. Specifically, we found that the Alphaproteobacteria-sister topology reported by Martijn et al. was the result of a very particular experimental setup and set of parameters that caused by loss of historical signal. In other cases of site excluded datasets, mitochondria emerged from within Alphaproteobacteria.

To detour the shortcomings of these methods, we here applied taxon replacement and selective exclusion in investigating the phylogenetic relationships between mitochondrial and alphaproteobacterial lineages. Supported by a number of bias-alleviated trees, we found that mitochondria have strong phylogenetic connection to the common ancestors of Rickettsiales and several fast-evolving alphaproteobacteria derived from marine surface metagenomes. While this result again supports a robust evolutionary association between mitochondria and Alphaproteobacteria, it also provides important ecological insights to the origin of both mitochondria and Rickettsiales. Based on our result, the common ancestor of mitochondria and Rickettsiales was a free-living alphaproteobacterium. This is consistent with a recent report favoring independent branching of Rickettsiales and mitochondria ^18^ but again in agreement with numerous previous studies which suggested phylogenetic connection between mitochondria and Rickettsiales ^6^.

Physiological and geological modellings have suggested that mitochondrial acquisition possibly occurred in shallow marine environments ^46^ or in anaerobic syntrophy ^47^. Our study along with others ^10^ implies that future work could discover the closest extant relatives of mitochondria in present marine environments. Proteome study of Rickettsiales and MarineAlpha bins in Alpha II may provide hints about the metabolic nature of the common ancestor of mitochondria ^47,48^.

## Methods

No statistical methods were used to predetermine sample size. The experiments were not randomized and the investigators were not blinded to allocation during experiments and outcome assessment.

### Implementation of site-exclusion metrics

To obtain the 24-alphamitoCOGs dataset in Martijn et al. (2018), file ‘alphaproteobacteria_mitochondria_untreated.aln’ was downloaded from https://datadryad.org//resource/doi:10.5061/dryad.068d0d0. As the names of some MarineAlpha bins in this file are not consistent with the phylogenetic trees in the original paper, we obtained the name mapping file from Dr. Joran Martijn on 4 July 2018. On this dataset, χ^2^-score based site exclusion was achieved by applying the equation introduced by Viklund et al. ^13^. ɀ-scores of sites were calculated according to the method introduced by Muñoz-Gómez et al. ^27^. Fast-evolving site exclusion was based on conditional mean site rates estimated under the LG+C60+F+R6 model in IQTREE (v1.5.5) using the ‘-wsr’ flag ^29^. Based on these three metrics, 5%, 10%, 20%, 40% and 60% of sites with the highest scores were excluded for downstream phylogenetic analyses. Moreover, site exclusion based on Stuart’s test was conducted by using the stationary-trimming function in BMGE (v1.12) ^32^.

Bowker’s test of symmetry ^31^ was used to produce subsets of the ‘alphamitoCOGs-24’ dataset in Martijn et al. (2018) by meeting increasingly stringent p-value-based thresholds (>0.005, >0.01, >0.05, >0.1, >0.2, >0.3, >0.4 and >0.5, respectively). The Bowker’s test has long been used as an overall test for symmetry ^31^. The test assesses symmetry in an r × r contingency table with the ij-th cell containing the observed frequency n_ij_. The null hypothesis for symmetry is H_0_ = n_ij_ = n_ji_, *i ≠ j, i,j = 1,…,r*, and the test value is computed as:

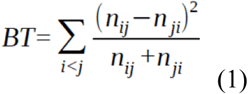

The test statistics follows *χ*^2^ distribution with the number of degrees of freedom equal to the number of comparisons (n_ij_ vs n_ji_) made.

The scoring function (SF) utilized for symmetry-based alignment trimming employed here is a sum of absolute values of natural logarithms of Bowker’s test’s p-values, each raised to a certain power (15 as the default value). SF can be computed as a mean over the values in an upper or lower triangular part of a square matrix which rows and columns represent taxa, populated with |ln *p*|^x^ values for Bowker’s tests among these taxa, e.g:

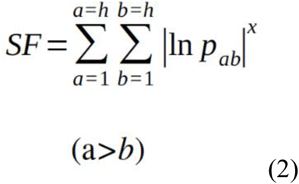

wherein *h* is the number of taxa in the msa, and p_ab_ is a p value for the sequences *a* and *b*.

The script which performs symmetry-based trimming (symmetry.pl, available as **Supplementary Data Files**) deletes a site in an alignment, computes a SF value and restores the original alignment. The operation is performed for every alignment site. Then, the site which removal results in lowest SF value is deleted irreversibly. The procedure is repeated for each shortened alignment subset until the lowest p-value for a pair-wise Bowker’s test in the trimmed dataset exceeds certain p-value-based threshold(s).

Exponentiation in formula 2 leads to a sooner recovery of trimmed subsets. The exponentiation disproportionally increases the addend values in formula 2 (|ln *p*_ab_|^x^) for smaller p values. For instance, the default addend in the formula 2 for p-value 0.5 is 0.004 and the addend for p-value 0.005 is 72789633288. Thus, when there is a disparity in individual p-values in the data, which is the case when the method is needed, the exponentiation increases the relative contribution of the lowest p-values onto the SF value size. At each trimming step the heuristic algorithm identifies a site which removal is likely to improve the worst (lowest) p-values. The script outputs a trimmed subset when the lowest p-value exceeds the threshold value. The suggested exponentiation, causing preferential improvement of the worst p-values at each site stripping step, is able to deliver a result when less positions are removed. The default exponent value (x = 15) has been determined experimentally.

### Phylogenetic inference and tree topology comparison

ML trees in this study were reconstructed by using IQTREE under either auto-selected simple model (ModelFinder) or mixed model (LG+C60+F) as specified in text. Bayesian trees were produced by using PhyloBayes MPI (v1.8) ^49^, four chains were run until a Maxdiff < 0.3 were reached.

For comparison of topology, ML trees of site-excluded datasets were first rooted to Beta-, Gammaproteobacteria, Magnetococcales, MarineProteo1 Bin1 and Bin 2. The dissimilarity value between each tree and the untreated tree was then calculated by using the Alignment metric developed by Nye et al. Briefly the Alignment metric considers all the ways that the branches of one tree map onto the other ^33^. The code was adapted from Kuhner et al. (2015) and implemented in Python.

### Genome and marker protein selection of the GC-bias-reduced dataset

The ‘18-alphamitoCOGs’ dataset of this study was based on the ‘24-alphamitoCOGs’ dataset in Martijn et al. (2018) after several modifications. Specifically, MAGs derived from composite bins, which contain sequences from multiple naturally existing genomes were excluded to minimize possible assembly-induced artifacts. Mitochondria and Rickettsiales in the original dataset used were replaced by less AT-rich alternatives (**Supplementary Table 2**). All relevant genomes were downloaded from the RefSeq database of NCBI on 21 July 2018.

For quality control of the 24 marker proteins of the original dataset, sequences of these proteins were downloaded from the MitoCOGs ^50^ database and then aligned by using MAFFT-L-INS-I (v 7.055b) ^51^, respectively. Alignment of each protein was trimmed by using trimAl (v.1.4) ^52^. Protein-specific e-values were determined with distributions of positive and negative sequences. For each gene, sequences classified into the proteins in MitoCOGs database were used as positive dataset and sequences classified into other proteins were used as negative one. E-value distribution of positive and negative sequences was calculated by using Hmmer (v3.2.1) ^53^. Protein-specific e-values were the minimum of 95% quantile e-values of positive sequences, and the minimum of negative sequences. We searched these 24 proteins individually in the genomes by using Hmmer based on protein-specific e-values of the HMM models. The obtained proteins were processed for ML tree reconstruction by using IQTREE under the model ‘LG+C60+F’. Copies identified as paralogs, possible contaminants or events of lateral gene transfer in each gene tree were removed. *Candidatus* Paracaedibacter symbiosus was excluded as multiple contaminant proteins were detected in its genome and we think its genome likely suffers from heavy contamination. MitoCOG0003 and MitoCOG0133 were excluded as they were detected in few genomes. MitoCOG00052, MitoCOG00060, MitoCOG00066 and MitoCOG00071 were excluded as they were absent in reselected mitochondrial genomes. Consequenly, 18 marker proteins were selected. Except for outgroup species (including Beta-, Gammaproteobacteria and Magnetococcales), genomes contained 16 or more than 16 of the 18 marker proteins were kept. Furthermore, we removed redundant MarineAlpha bins of the original dataset based on pairwise similarity of marker proteins by using BLASTP (v2.6.0+, identity ⩾ 0.99 and coverage ⩾ 0.95) to reduce computational time. As a result, 61 genomes were kept for downstream analysis.

Before phylogenetic inference, selected proteins were aligned respectively by using MAFFT-L-INS-i. Low-quality columns were removed by BMGE (-m BLOSUM30) and the multiple sequence alignments after quality control were concatenated.

## Acknowledgements

This work was financially supported by the National Natural Science Foundation of China (91851210, 41530105 and 81774152), the European Research Council (ERC 666053), the Shenzhen Key Laboratory of Marine Archaea Geo-Omics, Southern University of Science and Technology, (ZDSYS201802081843490), Shenzhen Science and Technology Innovation Commission (JCYJ20180305123458107), the VW foundation (93 046), and the Laboratory for Marine Geology, Qingdao National Laboratory for Marine Science and Technology, (MGQNLM-TD201810).

## Author Contributions

L.F., W.F.M. and R.Z. conceived this study. L.F., D.W., V.G., J.X., Y.X. and S.G. were involved in data analysis. L.F., V.G., C.Z, W.F.M. and R.Z. interpreted the results and drafted the manuscript. All authors participated in the critical revision of the manuscript.

## Competing interests

The authors declare no competing interests.

