## Supplementary tables and figures for "Mitochondria branch within Alphaproteobacteria"

**Supplementary Table 1 |**

| Site-exclusion methods | % of sites excluded | Alignment length | Datasets with all species | | | | | | |
|---|---|---|---|---|---|---|---|---|---|
| | | | Simple model | | | | Mixed model (C60) | | |
| | | | Model selected | Dissimilarity to the untreated tree (Align) | Mitochondria placement | Node bootstrap support | Dissimilarity to the untreated tree (Align) | mitochondria placement | Node bootstrap support |
| untreated | 0 | 6654 | LG+F+R8 | NA | IN | 79 | NA | IN | 95 |
| Stuart's test | 42.30% | 3840 | LG+F+R7 | 13.0324662 | OUT | 100 | 13.8870341 | OUT | 100 |
| Bowker's test (p>0.005) | 27.94% | 4795 | LG+F+R8 | 10.74363655 | IN | 92 | 13.2459772 | OUT | 100 |
| Bowker's test (p>0.01) | 29.07% | 4720 | LG+F+R8 | 10.74363655 | IN | 94 | 13.4844772 | OUT | 100 |
| Bowker's test (p>0.05) | 32.36% | 4501 | LG+F+R8 | 10.75818548 | IN | 47 | 13.0860239 | OUT | 100 |
| Bowker's test (p>0.1) | 34.66% | 4348 | LG+F+R8 | 9.134610879 | IN | 93 | 12.6144591 | OUT | 100 |
| Bowker's test (p>0.2) | 38.76% | 4075 | LG+F+R8 | 10.20127755 | IN | 94 | 12.3289127 | OUT | 100 |
| Bowker's test (p>0.3) | 42.09% | 3853 | LG+F+R8 | 10.29927218 | IN | 97 | 14.8906344 | OUT | 100 |
| Bowker's test (p>0.4) | 46.11% | 3586 | LG+F+R8 | 12.13805016 | IN | 97 | 13.7822456 | IN | 64 |
| Bowker's test (p>0.5) | 50.26% | 3310 | LG+F+R7 | 13.22756964 | IN | 90 | 14.8748392 | IN | 92 |
| $\chi$2-score | 5% | 6322 | LG+F+R8 | 3.54680197 | OUT | 100 | 4.62785046 | OUT | 100 |
| $\chi$2-score | 10% | 5989 | LG+F+R8 | 4.52903565 | OUT | 100 | 7.67034798 | OUT | 100 |
| $\chi$2-score | 20% | 5324 | LG+F+R8 | 7.86155725 | OUT | 100 | 10.3102749 | OUT | 100 |
| $\chi$2-score | 40% | 3993 | LG+F+R7 | 9.50723507 | OUT | 100 | 12.1129277 | OUT | 100 |
| $\chi$2-score | 60% | 2662 | LG+F+R7 | 12.3285779 | IN | 71 | 16.1858319 | OUT | 80 |
| $z$-score | 5% | 6321 | LG+F+R8 | 1.55192308 | IN | 89 | 1.42083333 | IN | 94 |
| $z$-score | 10% | 5989 | LG+F+R8 | 1.54358974 | IN | 90 | 1.76381579 | IN | 92 |
| $z$-score | 20% | 5323 | LG+F+R8 | 0.57142857 | IN | 69 | 3.58589744 | IN | 94 |
| $z$-score | 40% | 3993 | LG+F+R7 | 4.35548654 | IN | 90 | 2.56798246 | IN | 90 |

| | | | | | | | | | |
|---|---|---|---|---|---|---|---|---|---|
| ζ-score | 60% | 2659 | LG+F+R7 | 6.72778812 | IN | 89 | 5.15954402 | OUT | 100 |
| Fast-evolving | 5% | 6321 | LG+F+R7 | 1.32142857 | IN | 62 | 2.48237179 | IN | 56 |
| Fast-evolving | 10% | 5989 | LG+F+R7 | 3.04403131 | OUT | 100 | 3.04199866 | OUT | 100 |
| Fast-evolving | 20% | 5323 | LG+F+R6 | 3.11978022 | IN | 87 | 3.82566172 | IN | 74 |
| Fast-evolving | 40% | 3992 | LG+F+R4 | 6.55593328 | IN | 89 | 7.08341299 | IN | 92 |
| Fast-evolving | 60% | 2662 | LG+F+R3 | 18.0828063 | IN | 89 | 18.6806308 | OUT | 100 |

**Supplementary Table 2 | Taxa names, taxon group names, GC contents and amino acid ratios of the 18-alphamitoCOGs dataset.**

| Group | | | Taxa (abbreviation) | Order (NCBI) | Family (NCBI) | Genomic GC content | FYMINK/GARP ratio of marker proteins |
|---|---|---|---|---|---|---|---|
| outgroup | | | Pseudomonas aeruginosa PA7 | Pseudomonadales | Pseudomonadaceae | 0.665 | 0.877 |
| | | | Nitrosomonas sp. Is79A3 | Nitrosomonadales | Nitrosomonadaceae | 0.454 | 1.119 |
| | | | Chromobacterium violaceum ATCC 12472 | Neisseriales | Chromobacteriaceae | 0.644 | 0.960 |
| | | | Dechloromonas aromatica RCB | Rhodocyclales | Azonexaceae | 0.592 | 0.987 |
| others | | | Magnetococcus marinus MC-1 | Magnetococcales | Magnetococcaceae | 0.542 | 0.998 |
| backbone | Alpha I | Alpha Ia | Kordiimonas gwangyangensis | Kordiimonadales | Kordiimonadaceae | 0.575 | 0.982 |
| | | | alpha proteobacterium Q-1 | N.A. | N.A. | 0.561 | 0.914 |
| | | | Sphingomonas wittichii RW1 | Sphingomonadales | Sphingomonadaceae | 0.671 | 0.846 |
| | | | alpha proteobacterium JLT2015 | N.A. | N.A. | 0.641 | 0.874 |
| | | Alpha Ib | Parvibaculum lavamentivorans DS-1 | Rhizobiales | Rhodobiaceae | 0.623 | 0.915 |
| | | | Hyphomicrobium denitrificans ATCC 51888 | Rhizobiales | Hyphomicrobiaceae | 0.597 | 0.939 |
| | | | Ochrobactrum anthropi ATCC 49188 | Rhizobiales | Brucellaceae | 0.561 | 0.979 |
| | | | Caulobacter crescentus CB15 | Caulobacterales | Caulobacteraceae | 0.672 | 0.854 |
| | | | Parvularcula bermudensis HTCC2503 | Parvularculales | Parvularculaceae | 0.607 | 0.891 |
| | | | Rhodobacter sphaeroides 2.4.1 | Rhodobacterales | Rhodobacteraceae | 0.690 | 0.901 |
| | | | Roseobacter denitrificans OCh 114 | Rhodobacterales | Rhodobacteraceae | 0.589 | 0.956 |
| | Alpha II | Alpha IIa | MarineAlpha3 Bin2 (MA3B2) | N.A. | N.A. | 0.573 | 0.989 |
| | | | MarineAlpha3 Bin7 (MA3B7) | N.A. | N.A. | 0.395 | 1.143 |
| | | | Thalassospira xiamenensis | Rhodospirillales | Rhodospirillaceae | 0.548 | 0.976 |
| | | | Magnetospirillum magneticum AMB-1 | Rhodospirillales | Rhodospirillaceae | 0.651 | 0.931 |
| | | | Rhodospirillum rubrum ATCC 11170 | Rhodospirillales | Rhodospirillaceae | 0.654 | 0.923 |
| | | Alpha IIb | MarineAlpha9 Bin5 (MA9B5) | N.A. | N.A. | 0.501 | 0.959 |
| | | | MarineAlpha11 Bin1 (MA11B1) | N.A. | N.A. | 0.503 | 0.965 |
| | | | MarineAlpha12 Bin1 (MA12B1) | N.A. | N.A. | 0.398 | 1.077 |
| | Alpha III | Alpha IIIa | Kiloniella sp. P1-1 | Kiloniellales | Kiloniellaceae | 0.461 | 1.006 |
| | | | Candidatus Puniceispirillum marinum IMCC1322 | SAR116 cluster | N.A. | 0.489 | 0.977 |
| | | | Candidatus Endolissoclinum faulkneri | Rhodospirillales | Rhodospirillaceae | 0.341 | 1.220 |
| | | | alpha proteobacterium BAL199 | N.A. | N.A. | 0.650 | 0.880 |
| | | | Reyranella massiliensis | Rhodospirillales | N.A. | 0.646 | 0.886 |

| | | | | | | | |
|---|---|---|---|---|---|---|---|
| | | Alpha IIIb | Micavibrio aeruginosavorus ARL-13 | N.A. | N.A. | 0.550 | 0.964 |
| | | | Azospirillum brasilense Sp245 | Rhodospirillales | Rhodospirillaceae | 0.690 | 0.919 |
| | | | Rhodospirillum centenum SW | Rhodospirillales | Rhodospirillaceae | 0.705 | 0.881 |
| | | | Roseomonas cervicalis | Rhodospirillales | Acetobacteraceae | 0.627 | 0.831 |
| | | | Acidiphilium cryptum JF-5 | Rhodospirillales | Acetobacteraceae | 0.671 | 0.845 |
| | | | Gluconacetobacter hansenii ATCC 23769 | Rhodospirillales | Acetobacteraceae | 0.596 | 0.835 |
| | GT | | Geminicoccus roseus | Rhodospirillales | Geminicoccaceae | 0.685 | 0.873 |
| | | | Tistrella mobilis KA081020-065 | Rhodospirillales | Rhodospirillaceae | 0.684 | 0.845 |
| fast-evolving lineages | Holosporales | | Candidatus Caedibacter acanthamoebae / Paracaedimonas | Holosporales | Caedimonadaceae | 0.382 | 1.236 |
| | | | Candidatus Odyssella thessalonicensis | Holosporales | Candidatus Paracaedibacteraceae | 0.420 | 1.108 |
| | Rickettsiales | | Neorickettsia risticii str. Illinois | Rickettsiales | Anaplasmataceae | 0.413 | 1.264 |
| | | | Neorickettsia sp. 179522 | Rickettsiales | Anaplasmataceae | 0.411 | 1.281 |
| | | | Wolbachia pipientis | Rickettsiales | Wolbachieae | 0.382 | 1.484 |
| | | | Anaplasma marginale str. Florida | Rickettsiales | Anaplasmataceae | 0.498 | 0.982 |
| | | | Anaplasma ovis | Rickettsiales | Anaplasmataceae | 0.497 | 0.989 |
| | Pelagibacterales | | Candidatus Pelagibacter sp. IMCC9063 | Pelagibacterales | Pelagibacteraceae | 0.317 | 1.528 |
| | | | Candidatus Pelagibacter ubique HTCC1062 | Pelagibacterales | Pelagibacteraceae | 0.297 | 1.560 |
| | HIMB59 | | alphaproteobacterium HIMB59 | N.A. | N.A. | 0.323 | 1.646 |
| | FEMAG I | | MarineAlpha9 Bin4 (MA9B4) | N.A. | N.A. | 0.288 | 1.525 |
| | | | MarineAlpha9 Bin1 (MA9B1) | N.A. | N.A. | 0.322 | 1.336 |
| | | | MarineAlpha9 Bin3 (MA9B3) | N.A. | N.A. | 0.293 | 1.264 |
| | FEMAG II | | MarineAlpha6 Bin5 (MA6B5) | N.A. | N.A. | 0.257 | 1.834 |
| | | | MarineAlpha5 Bin12 (MA5B12) | N.A. | N.A. | 0.279 | 1.717 |
| | | | MarineAlpha5 Bin9 (MA5B9) | N.A. | N.A. | 0.270 | 1.748 |
| | | | MarineAlpha5 Bin7 (MA5B7) | N.A. | N.A. | 0.281 | 1.634 |
| | | | MarineAlpha5 Bin3 (MA5B3) | N.A. | N.A. | 0.310 | 1.568 |
| | | | MarineAlpha5 Bin5 (MA5B5) | N.A. | N.A. | 0.283 | 1.604 |
| | Mitochondria | | Calypogeia arguta mt | N.A. | N.A. | 0.456 | 1.273 |
| | | | Pleurozia purpurea mt | N.A. | N.A. | 0.454 | 1.329 |
| | | | Ophioglossum californicum mt | N.A. | N.A. | 0.522 | 0.955 |
| | | | Nelumbo nucifera mt | N.A. | N.A. | 0.482 | 1.135 |
| | | | Nymphaea colorata mt | N.A. | N.A. | 0.451 | 1.114 |

**Supplementary Table 3 | COGs used in the 18-alphamitoCOGs dataset.**

| COG ID | Protein annotation |
|---|---|
| MitoCOG0001 | NADH dehydrogenase subunit 2 |
| MitoCOG0004 | ATP synthase subunit 6 |
| MitoCOG0005 | cytochrome c oxidase subunit 3 |
| MitoCOG0008 | NADH dehydrogenase subunit 4 |
| MitoCOG0009 | NADH dehydrogenase subunit 5 |
| MitoCOG0010 | NADH dehydrogenase subunit 6 |
| MitoCOG0011 | cytochrome b |
| MitoCOG0012 | NADH dehydrogenase subunit 1 |
| MitoCOG0027 | ribosomal protein L2 |
| MitoCOG0030 | ribosomal protein S12 |
| MitoCOG0031 | NADH dehydrogenase subunit 7 |
| MitoCOG0039 | ribosomal protein L16 |
| MitoCOG0040 | cytochrome c maturation protein ccmFN |
| MitoCOG0043 | NADH dehydrogenase subunit 9 |
| MitoCOG0053 | ribosomal protein L6 |
| MitoCOG0055 | ribosomal protein S14 |
| MitoCOG0059 | ATP synthase subunit 1 |
| MitoCOG0067 | ribosomal protein S4 |

**Supplementary Figures**

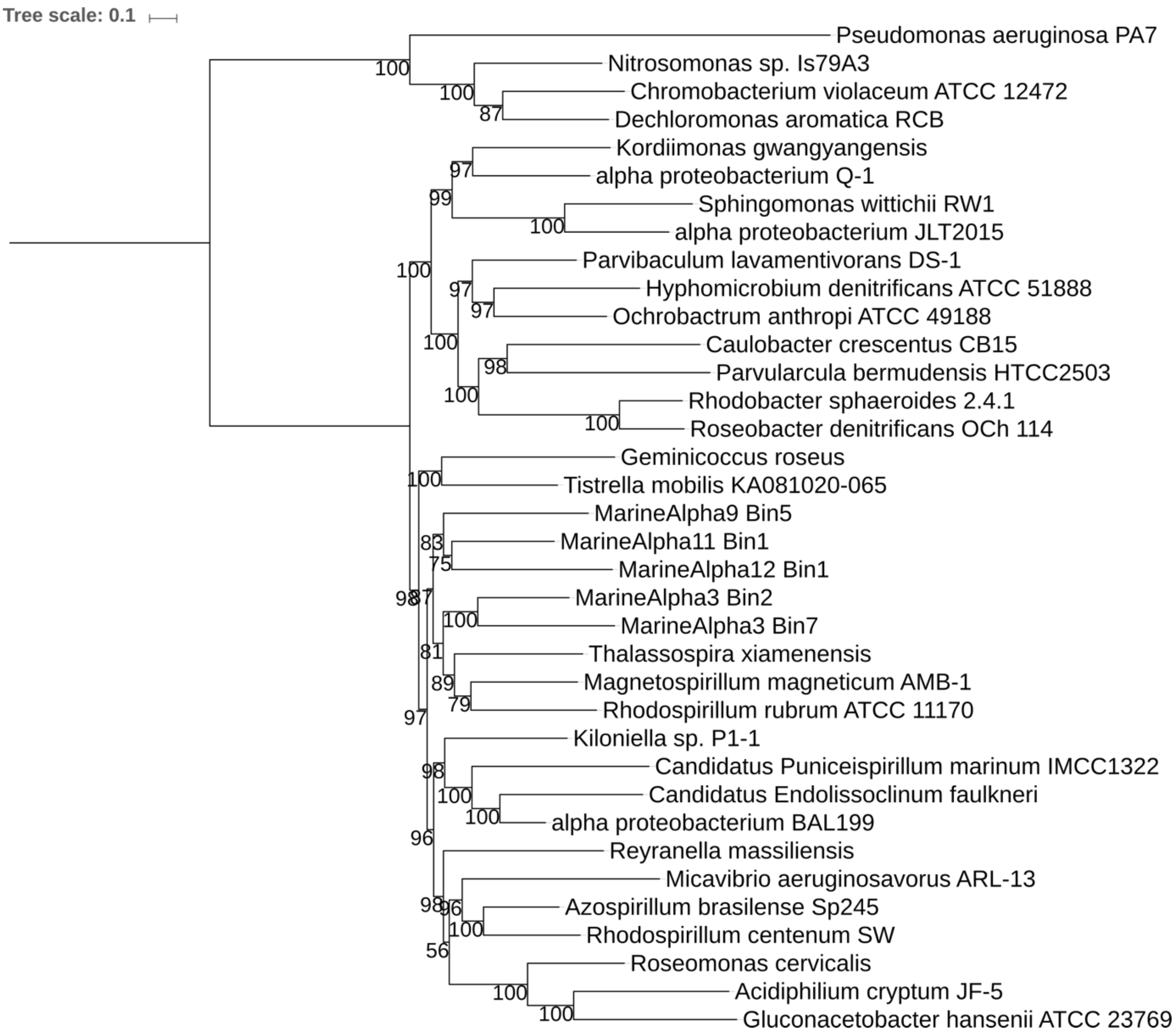


**Supplementary Fig. 1 | ML phylogenetic tree of backbone alphaproteobacteria in the 18-alphamitoCOGs dataset.** The tree is rooted with representatives of Beta- and Gammaproteobacteria. Node support values are based on the bootstrap results after 1000 iterations.

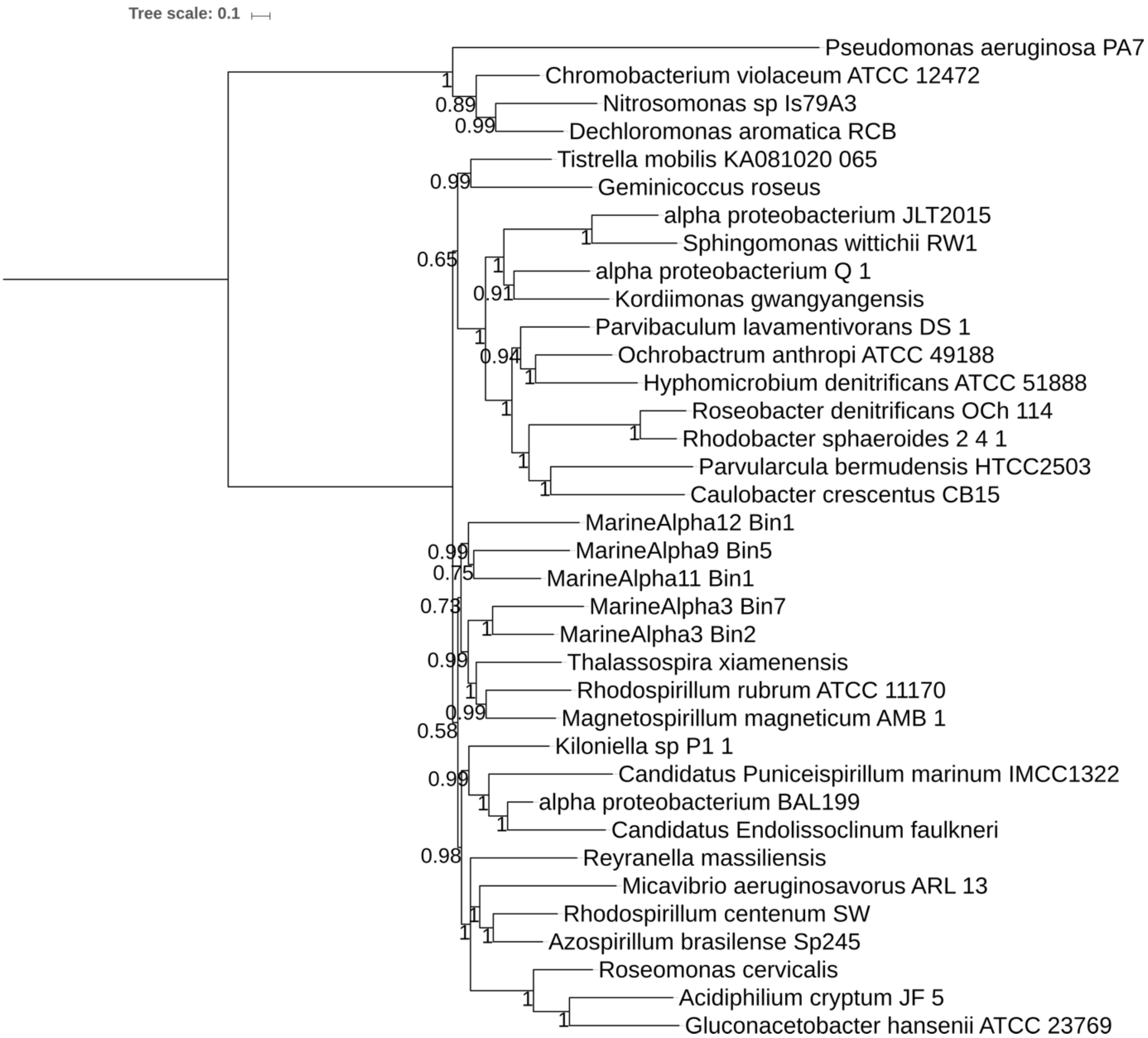


**Supplementary Fig. 2 | Bayesian phylogenetic tree of backbone alphaproteobacteria in the 18-alphamitoCOGs dataset.** The tree is rooted with representatives of Beta- and Gammaproteobacteria. Node values show posterior probability support values.

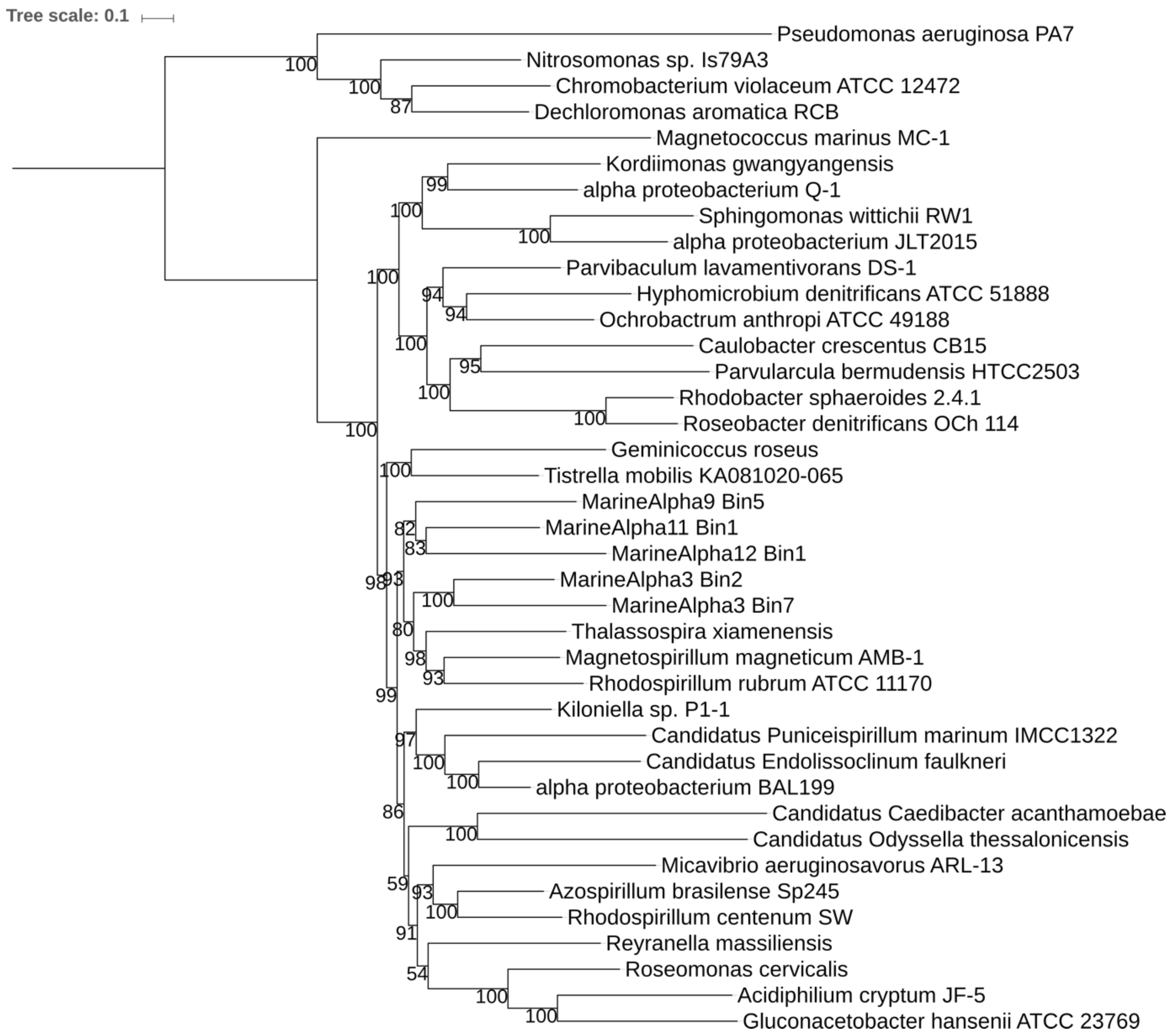


**Supplementary Fig. 3 | ML phylogenetic tree of backbone alphaproteobacteria and Holosporales in the 18-alphamitoCOGs dataset.** The tree is rooted with representatives of Beta- and Gammaproteobacteria. Node support values are based on the bootstrap results after 1000 iterations.

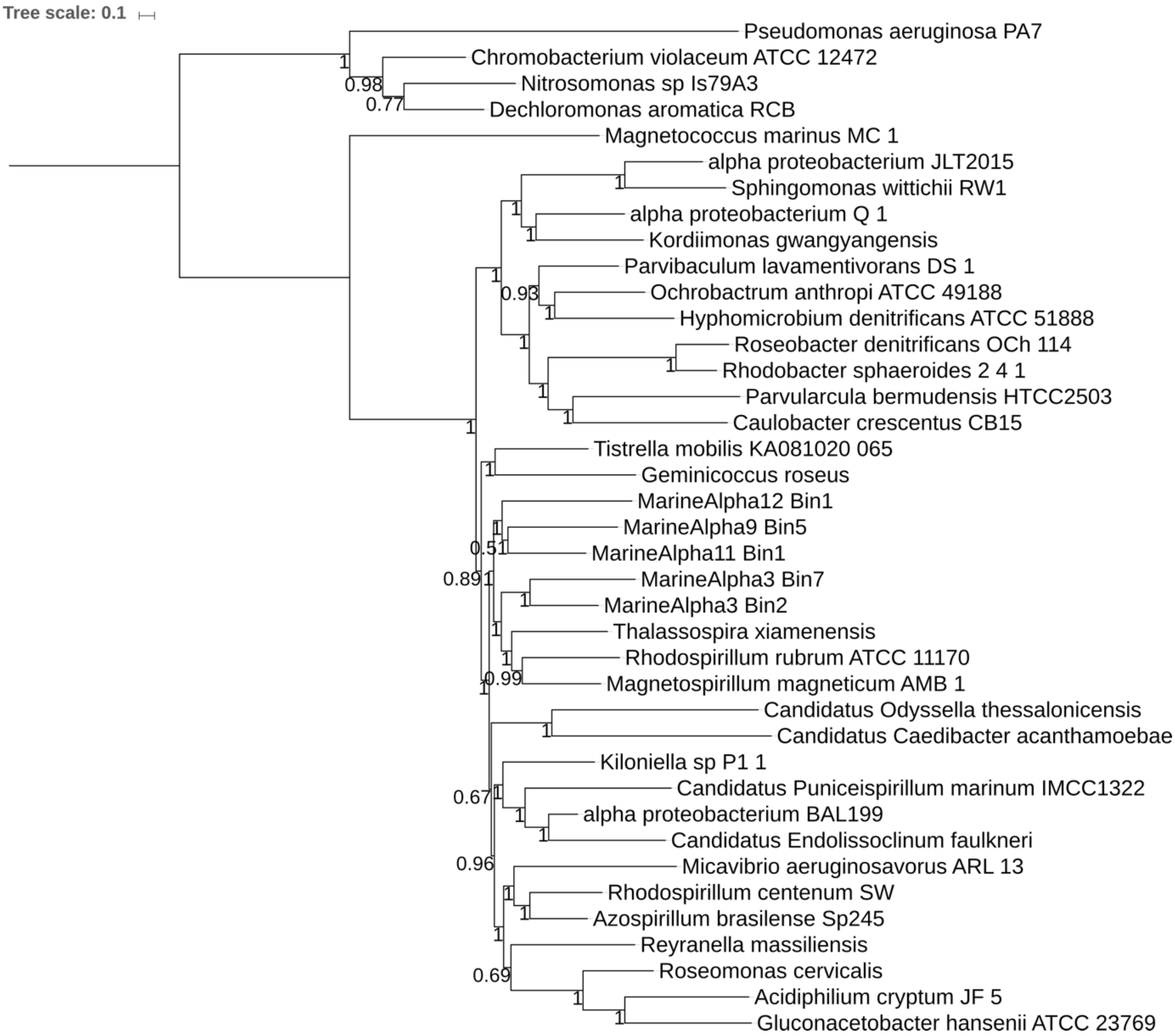


**Supplementary Fig. 4 | Bayesian phylogenetic tree of backbone alphaproteobacteria and Holosporales in the 18-alphamitoCOGs dataset.** The tree is rooted with representatives of Beta- and Gammaproteobacteria. Node values show posterior probability support values.

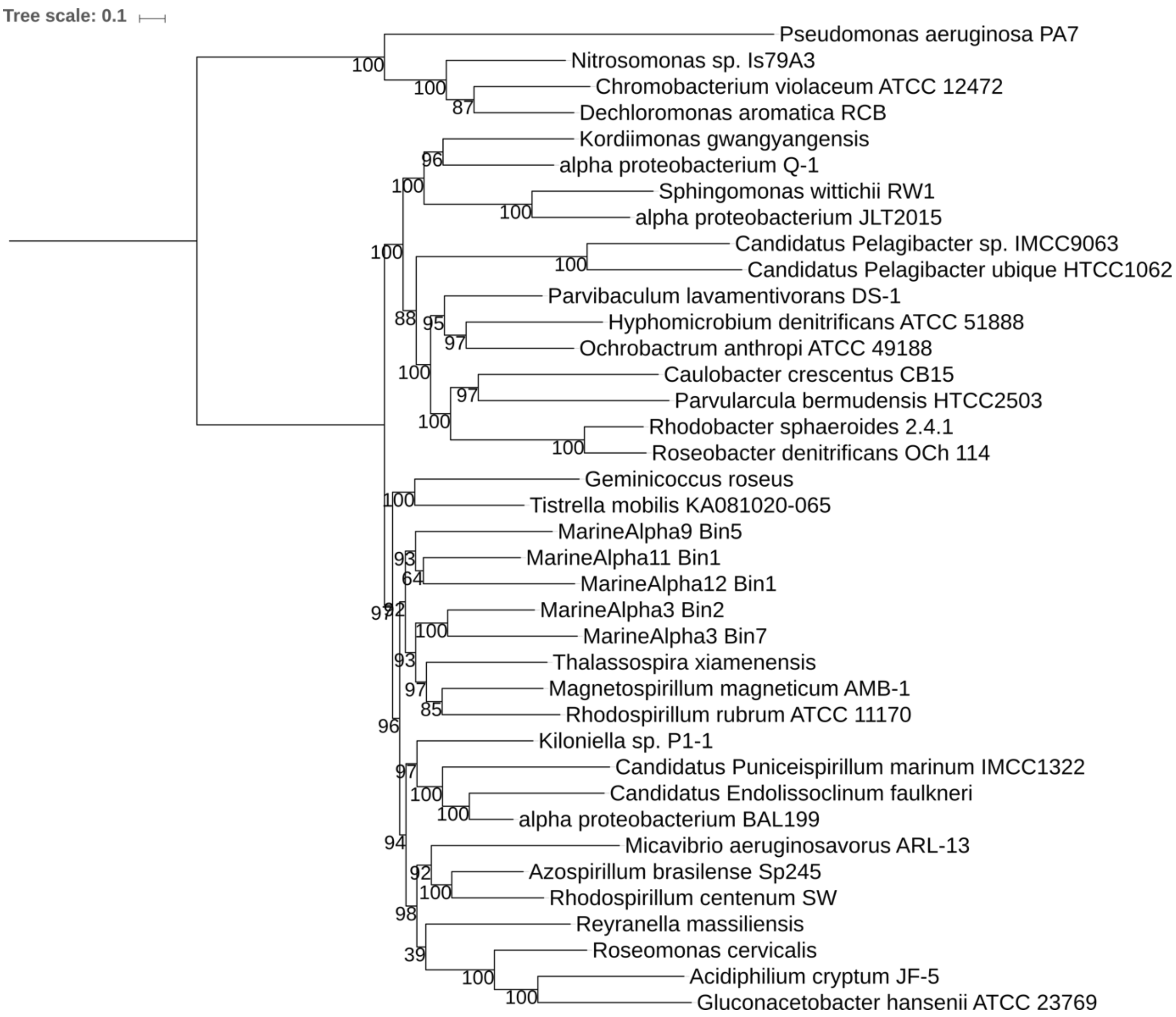


**Supplementary Fig. 5 | ML phylogenetic tree of backbone alphaproteobacteria and Pelagibacterales in the 18-alphamitoCOGs dataset.** The tree is rooted with representatives of Beta- and Gammaproteobacteria. Node support values are based on the bootstrap results after 1000 iterations.

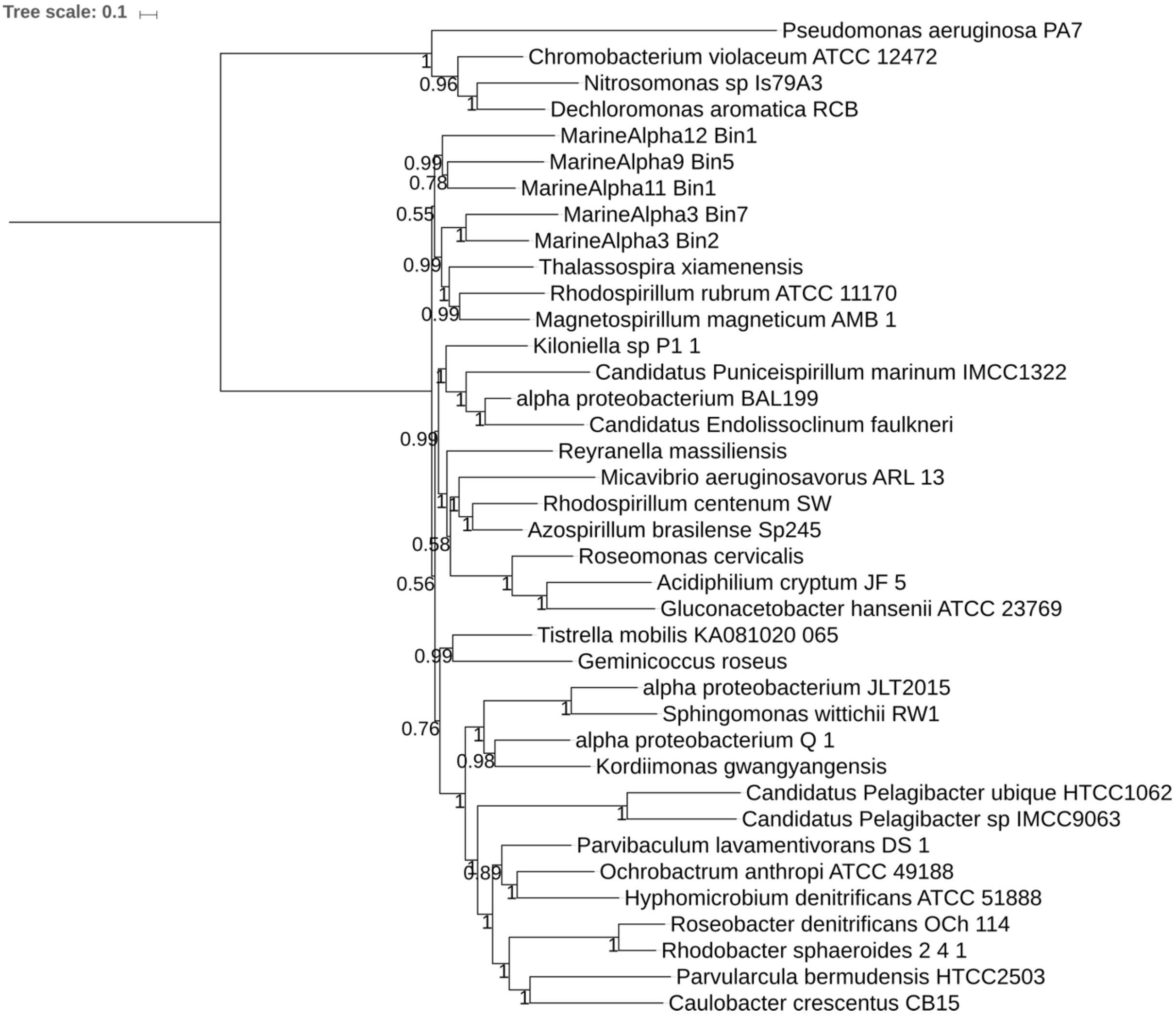


**Supplementary Fig. 6 | Bayesian phylogenetic tree of backbone alphaproteobacteria and Pelagibacterales in the 18-alphamitoCOGs dataset.** The tree is rooted with representatives of Beta- and Gammaproteobacteria. Node values show posterior probability support values.

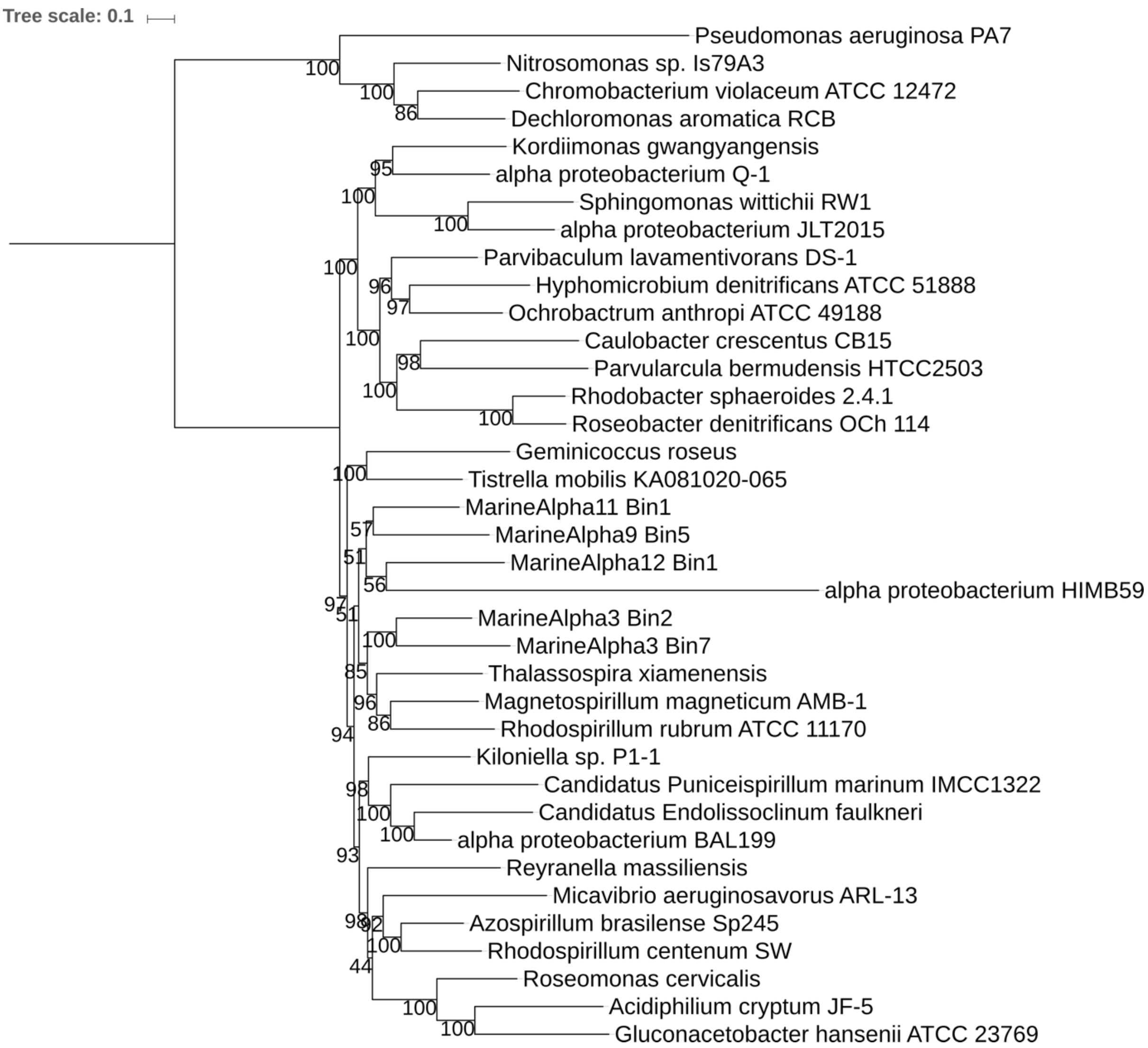


**Supplementary Fig. 7 | ML phylogenetic tree of backbone alphaproteobacteria and alphaproteobacterium HIMB59 in the 18-alphamitoCOGs dataset.** The tree is rooted with representatives of Beta- and Gammaproteobacteria. Node support values are based on the bootstrap results after 1000 iterations.

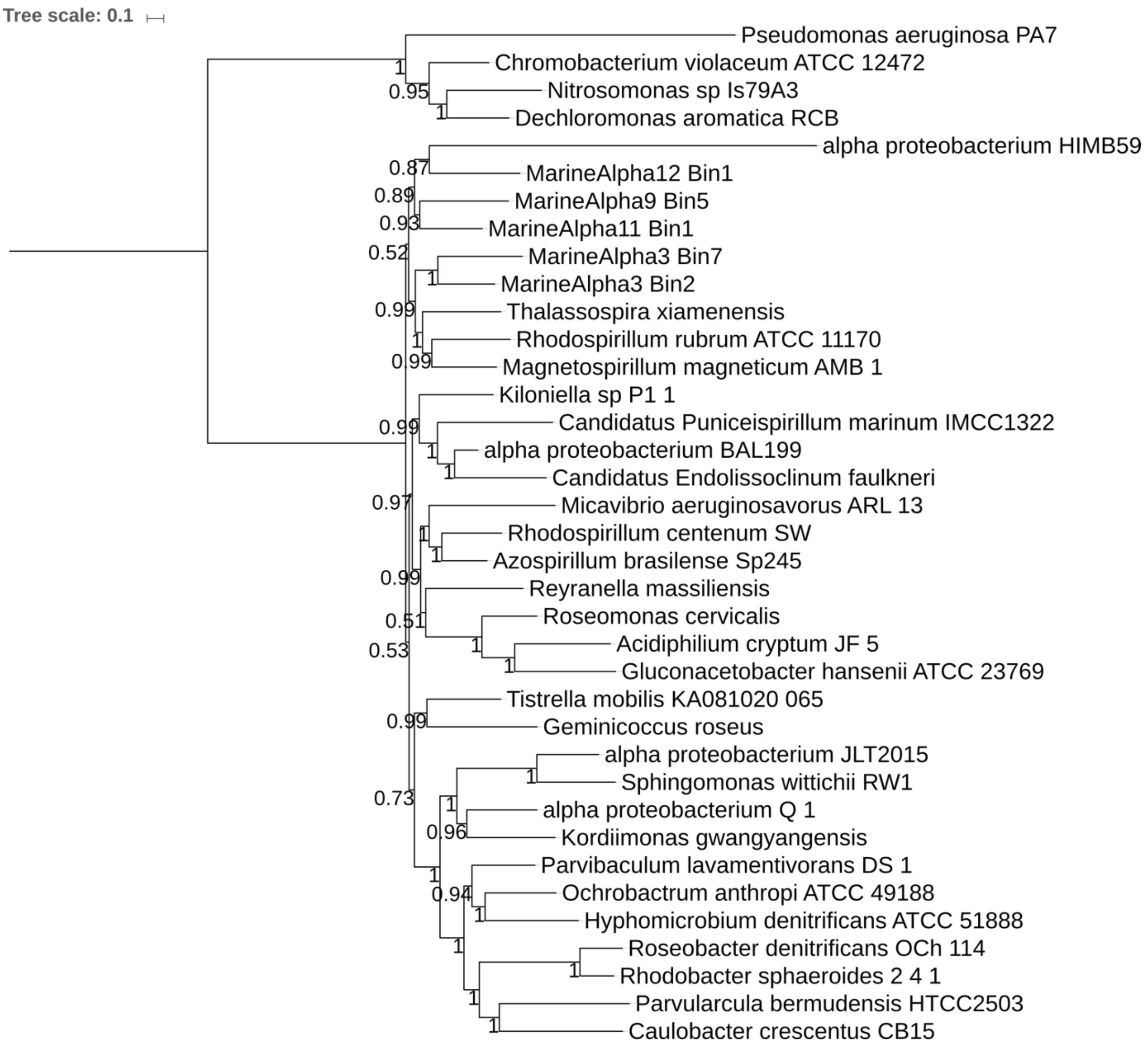


**Supplementary Fig. 8 | Bayesian phylogenetic tree of backbone alphaproteobacteria and alphaproteobacterium HIMB59 in the 18-alphamitoCOGs dataset.** The tree is rooted with representatives of Beta- and Gammaproteobacteria. Node values show posterior probability support values.

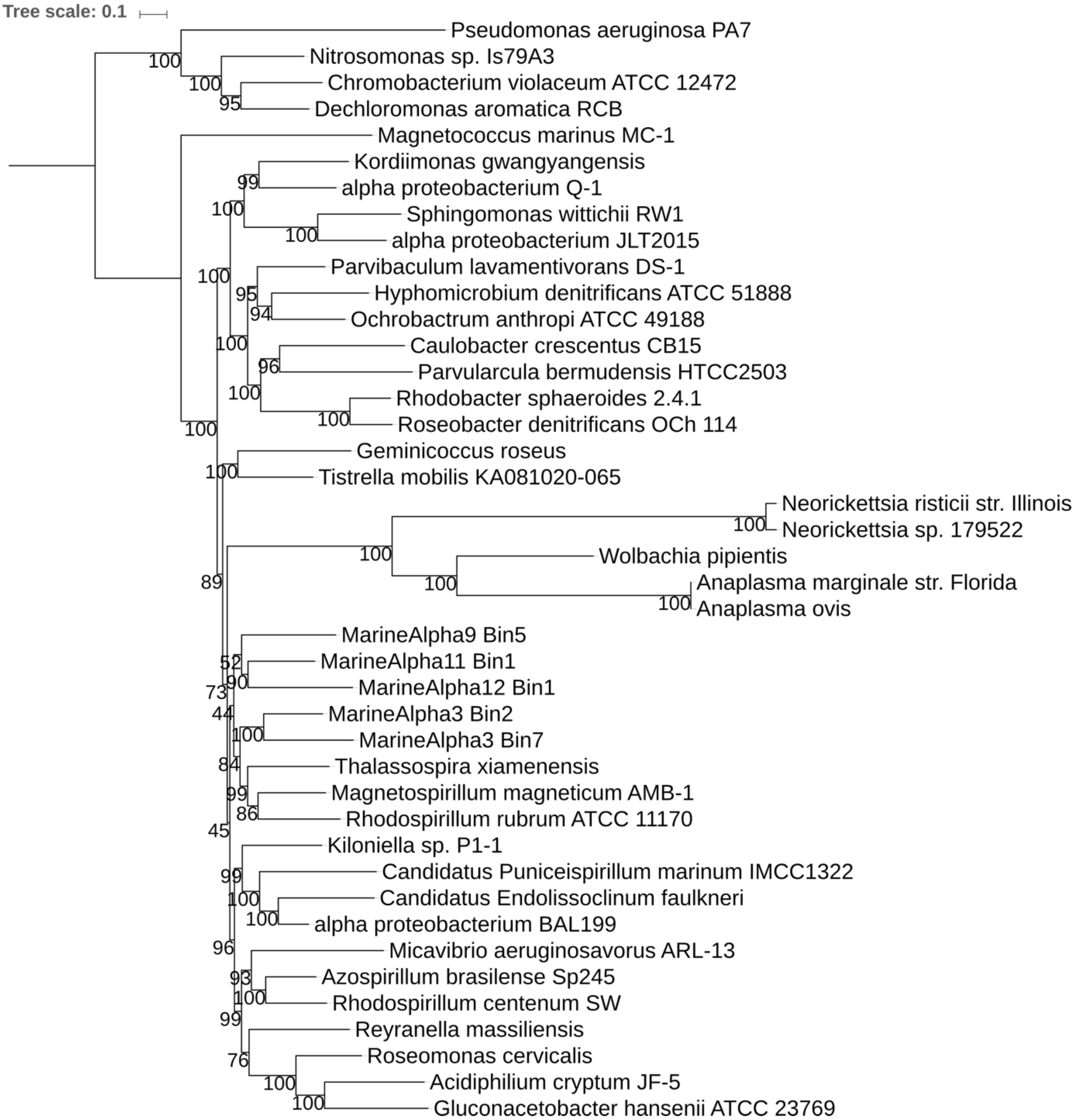


**Supplementary Fig. 9 | ML phylogenetic tree of backbone alphaproteobacteria and Rickettsiales in the 18-alphamitoCOGs dataset.** The tree is rooted with representatives of Beta- and Gammaproteobacteria. Node support values are based on the bootstrap results after 1000 iterations.

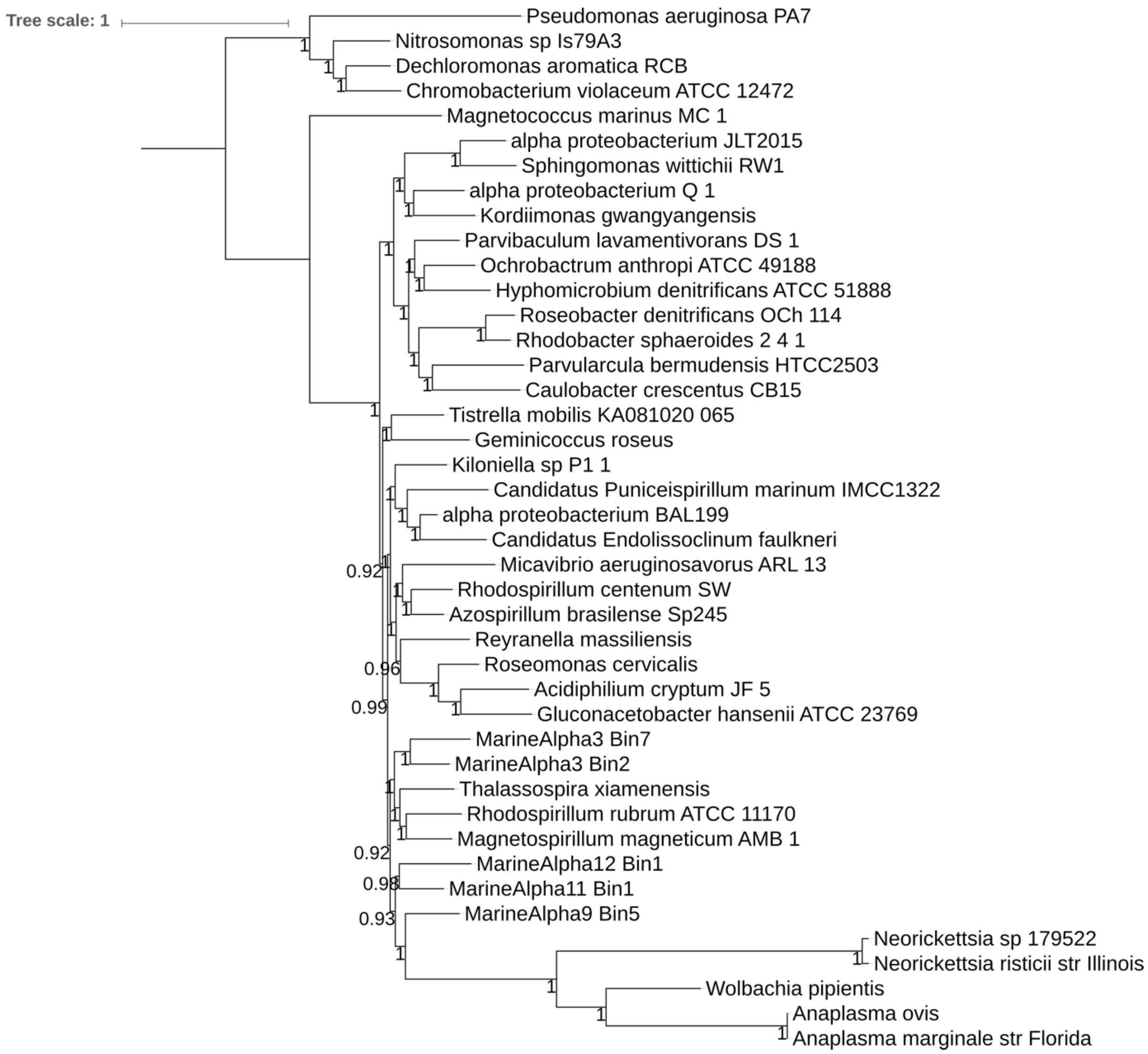


**Supplementary Fig. 10 | Bayesian phylogenetic tree of backbone alphaproteobacteria and Rickettsiales in the 18-alphamitoCOGs dataset.** The tree is rooted with representatives of Beta- and Gammaproteobacteria. Node values show posterior probability support values.

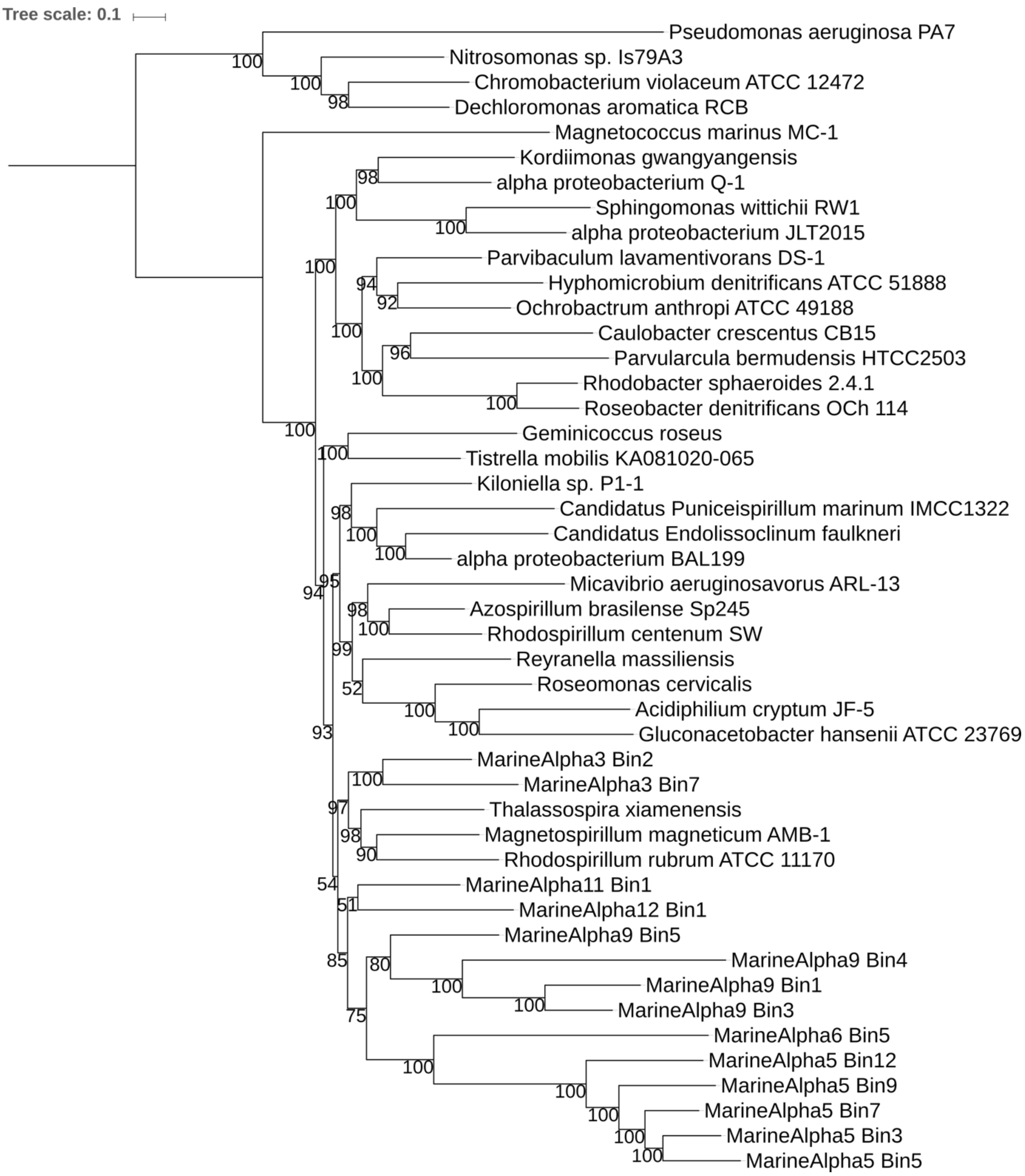


**Supplementary Fig. 11 | ML phylogenetic tree of backbone alphaproteobacteria, FEMAG I and FEMAG II in the 18-alphamitoCOGs dataset.** The tree is rooted with representatives of Beta- and Gammaproteobacteria. Node support values are based on the bootstrap results after 1000 iterations.

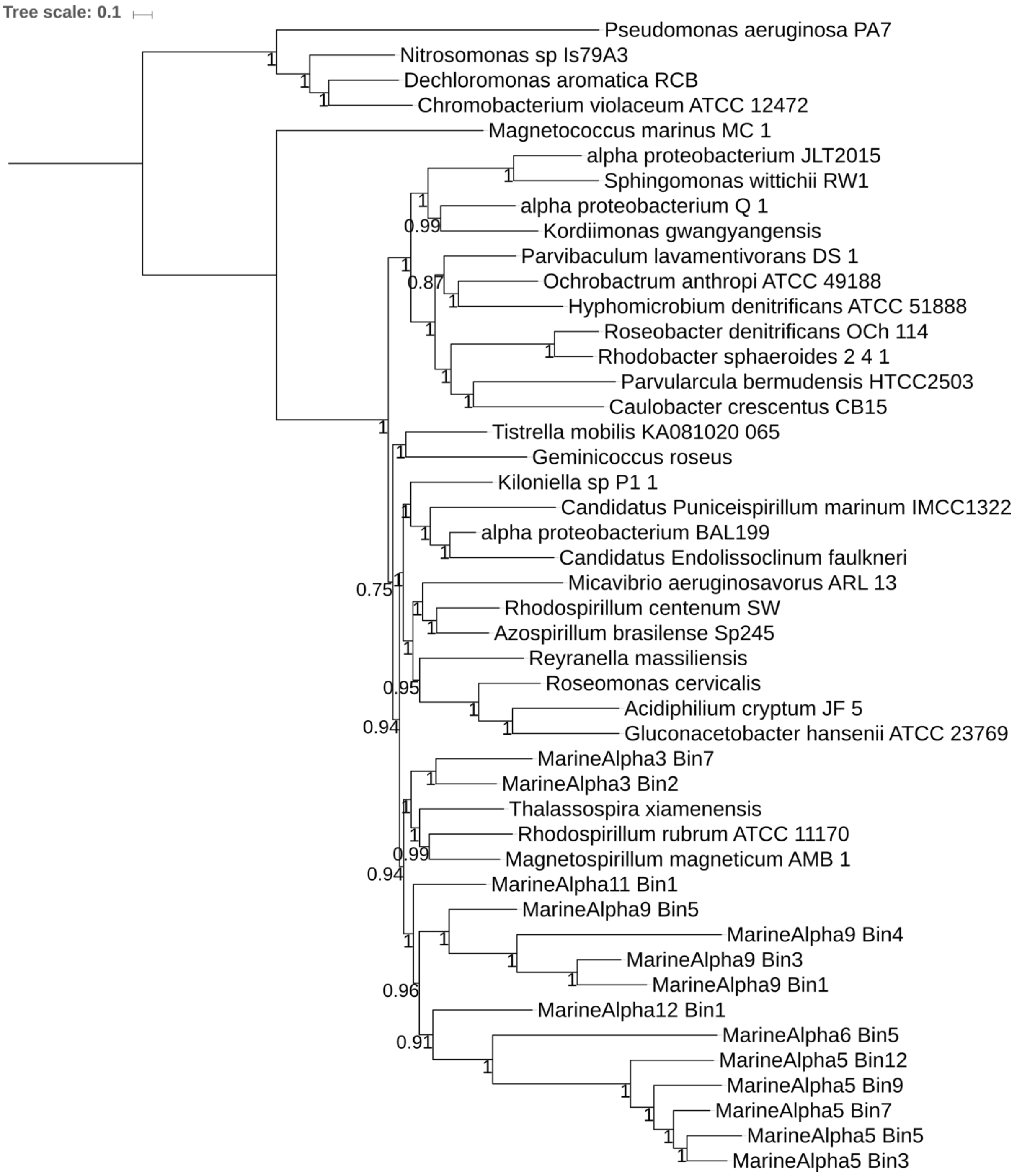


**Supplementary Fig. 12 | Bayesian phylogenetic tree of backbone alphaproteobacteria, FEMAG I and FEMAG II in the 18-alphamitoCOGs dataset.** The tree is rooted with representatives of Beta- and Gammaproteobacteria. Node values show posterior probability support values.

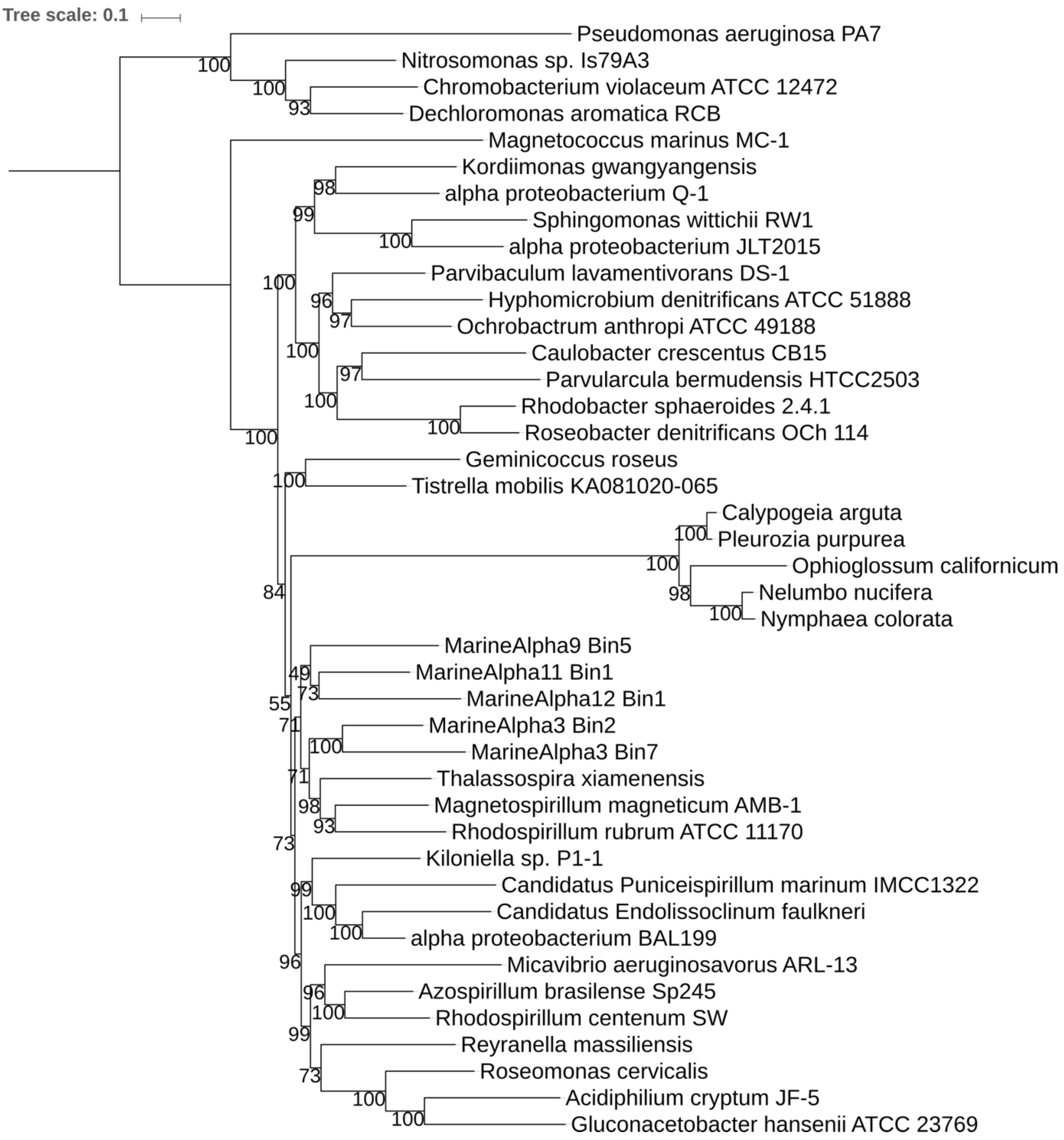


**Supplementary Fig. 13 | ML phylogenetic tree of backbone alphaproteobacteria and mitochondria in the 18-alphamitoCOGs dataset.** The tree is rooted with representatives of Beta- and Gammaproteobacteria. Node support values are based on the bootstrap results after 1000 iterations.

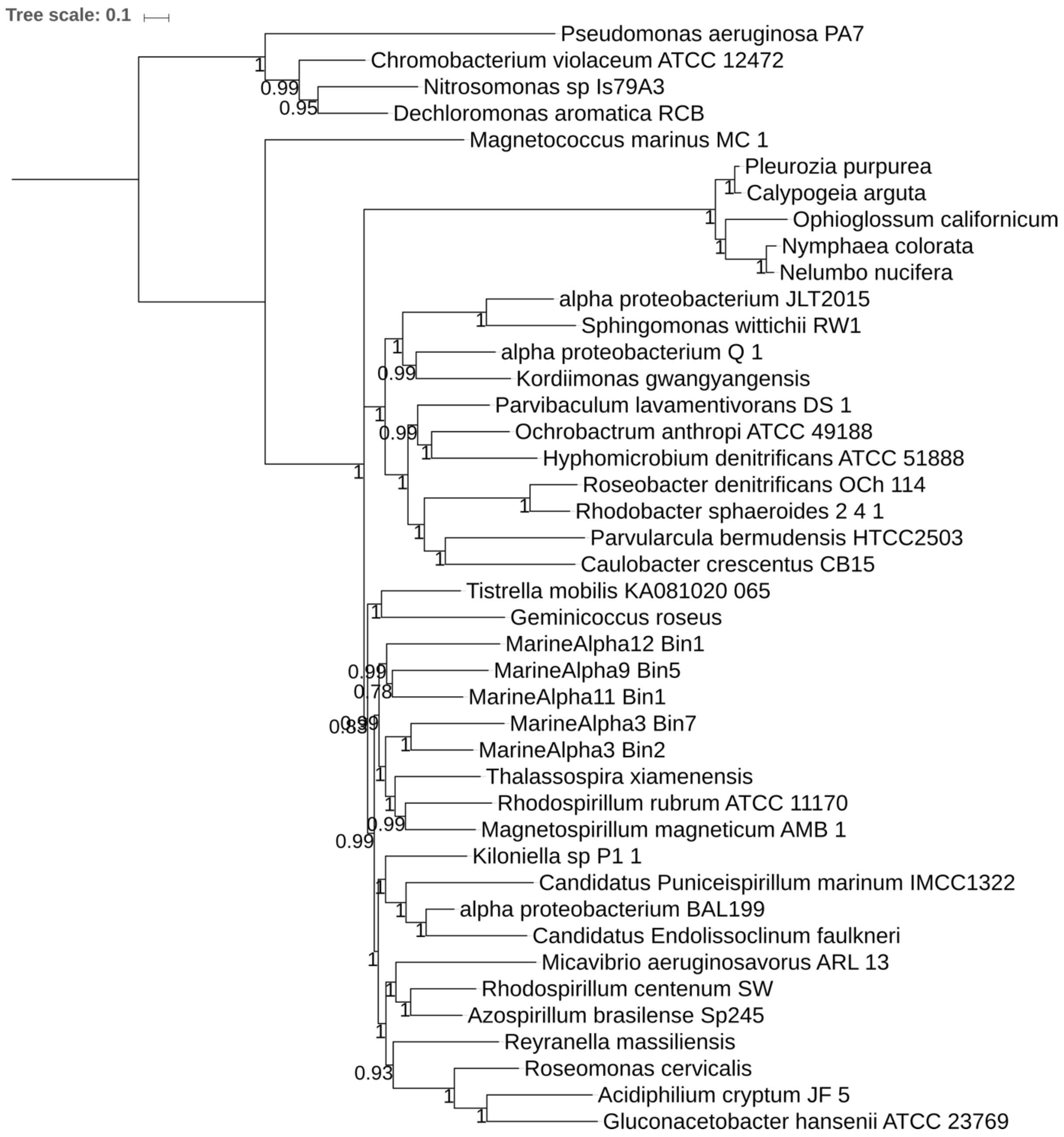


**Supplementary Fig. 14 | Bayesian phylogenetic tree of backbone alphaproteobacteria and mitochondria in the 18-alphamitoCOGs dataset.** The tree is rooted with representatives of Beta- and Gammaproteobacteria. Node values show posterior probability support values.

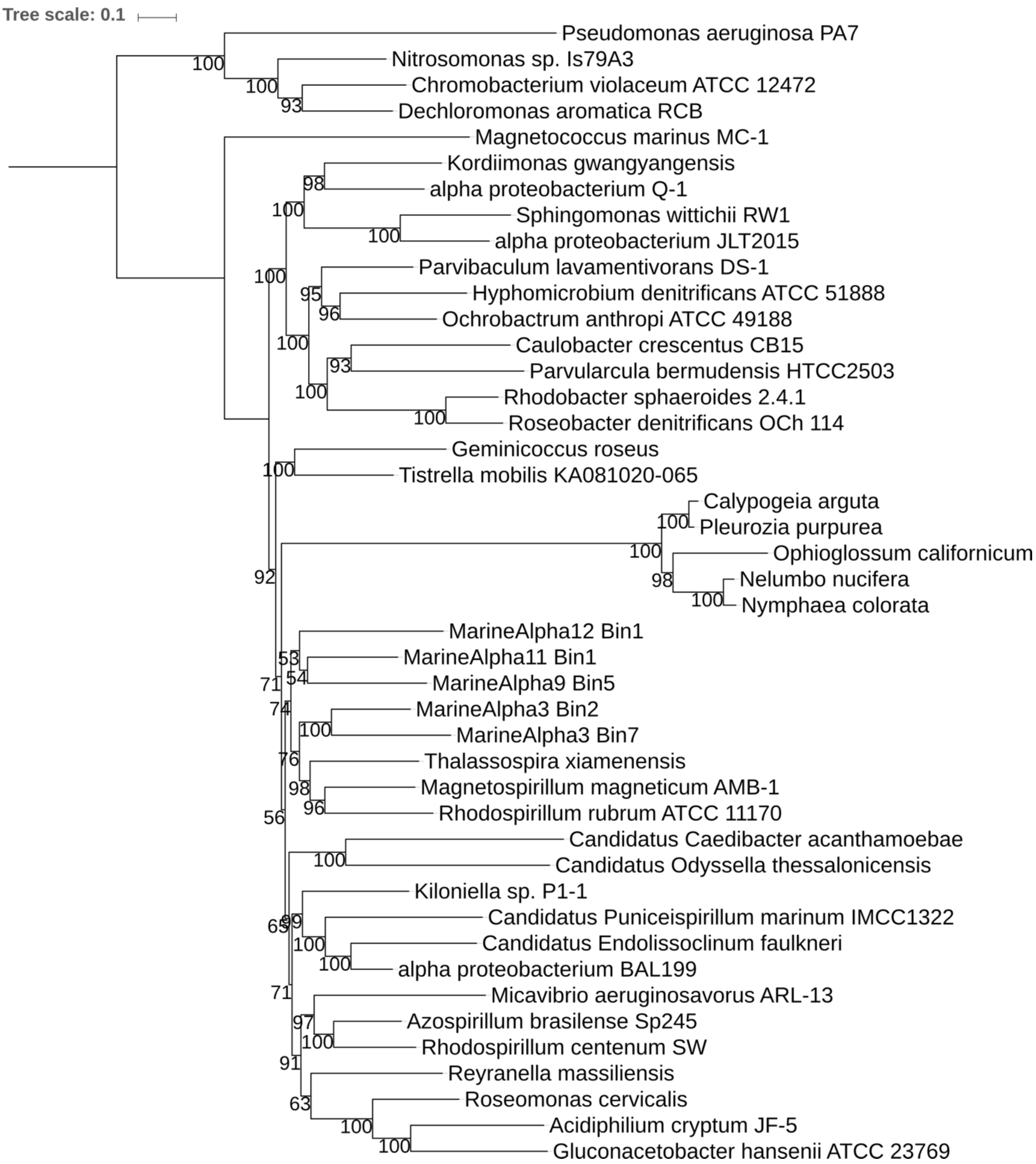


**Supplementary Fig. 15 | ML phylogenetic tree of backbone alphaproteobacteria, Holosporales and mitochondria in the 18-alphamitoCOGs dataset.** The tree is rooted with representatives of Beta- and Gammaproteobacteria. Node support values are based on the bootstrap results after 1000 iterations.

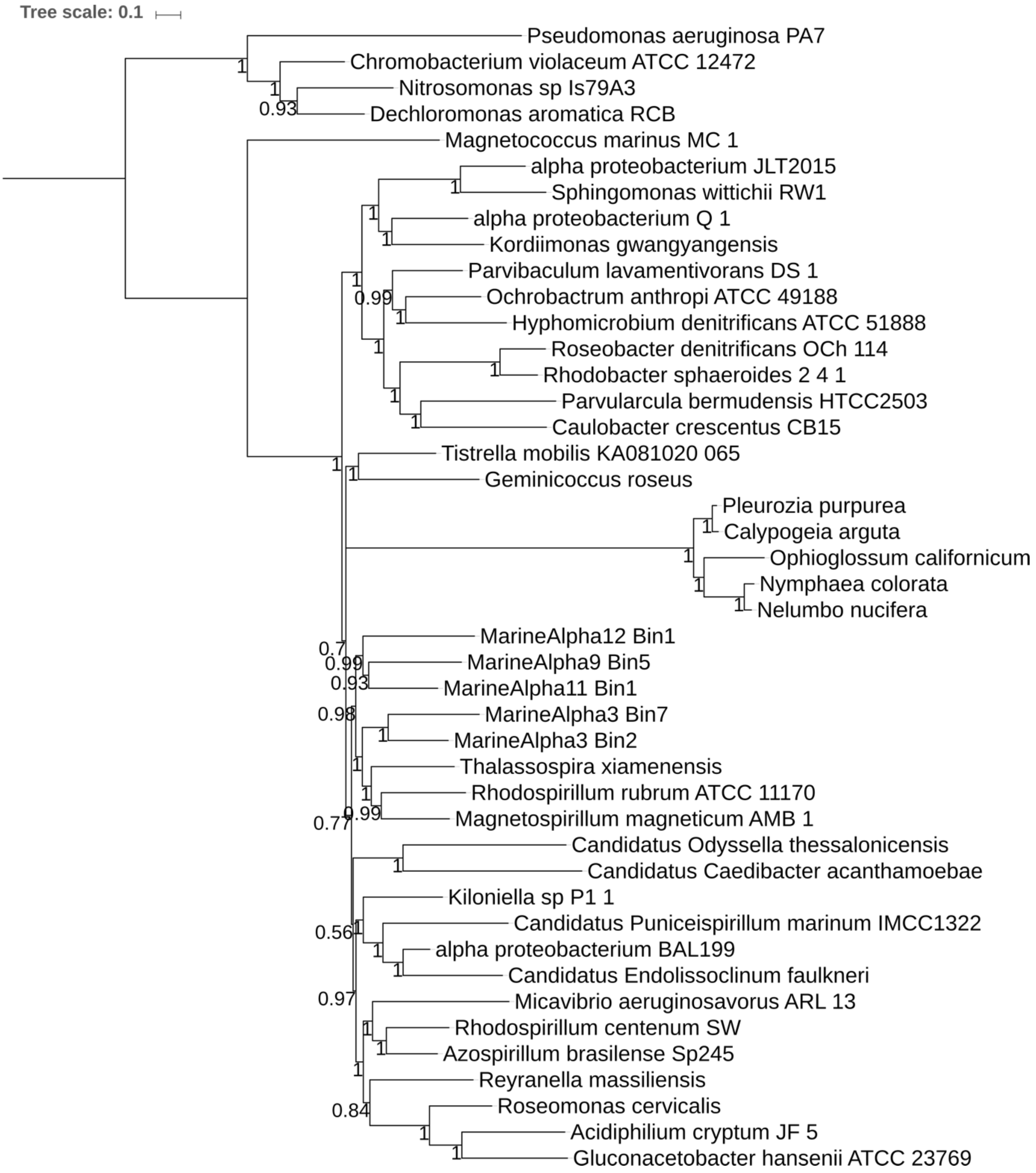


**Supplementary Fig. 16 | Bayesian phylogenetic tree of backbone alphaproteobacteria, Holosporales and mitochondria in the 18-alphamitoCOGs dataset.** The tree is rooted with representatives of Beta- and Gammaproteobacteria. Node values show posterior probability support values.

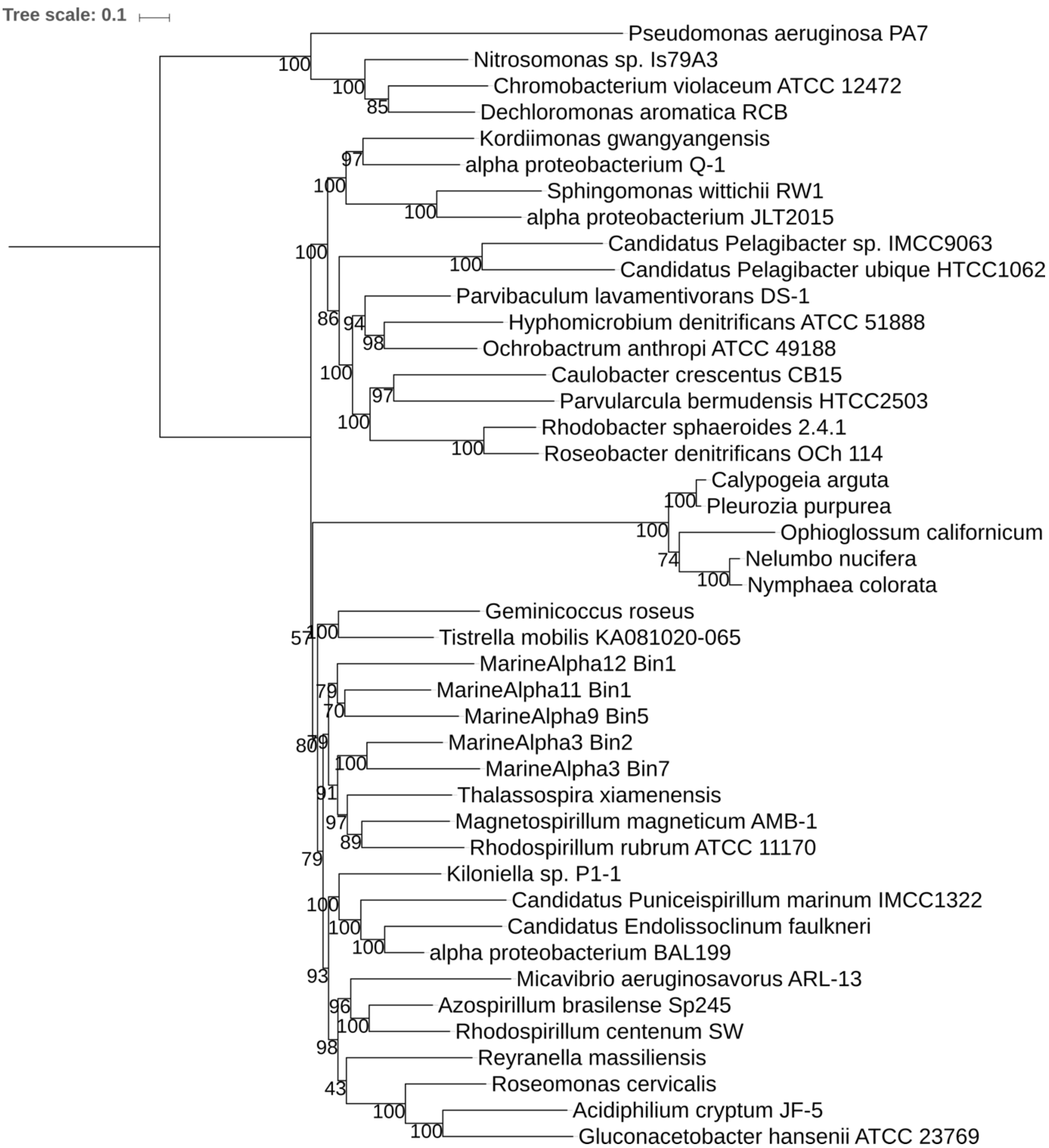


**Supplementary Fig. 17 | ML phylogenetic tree of backbone alphaproteobacteria, Pelagibacterales and mitochondria in the 18-alphamitoCOGs dataset.** The tree is rooted with representatives of Beta- and Gammaproteobacteria. Node support values are based on the bootstrap results after 1000 iterations.

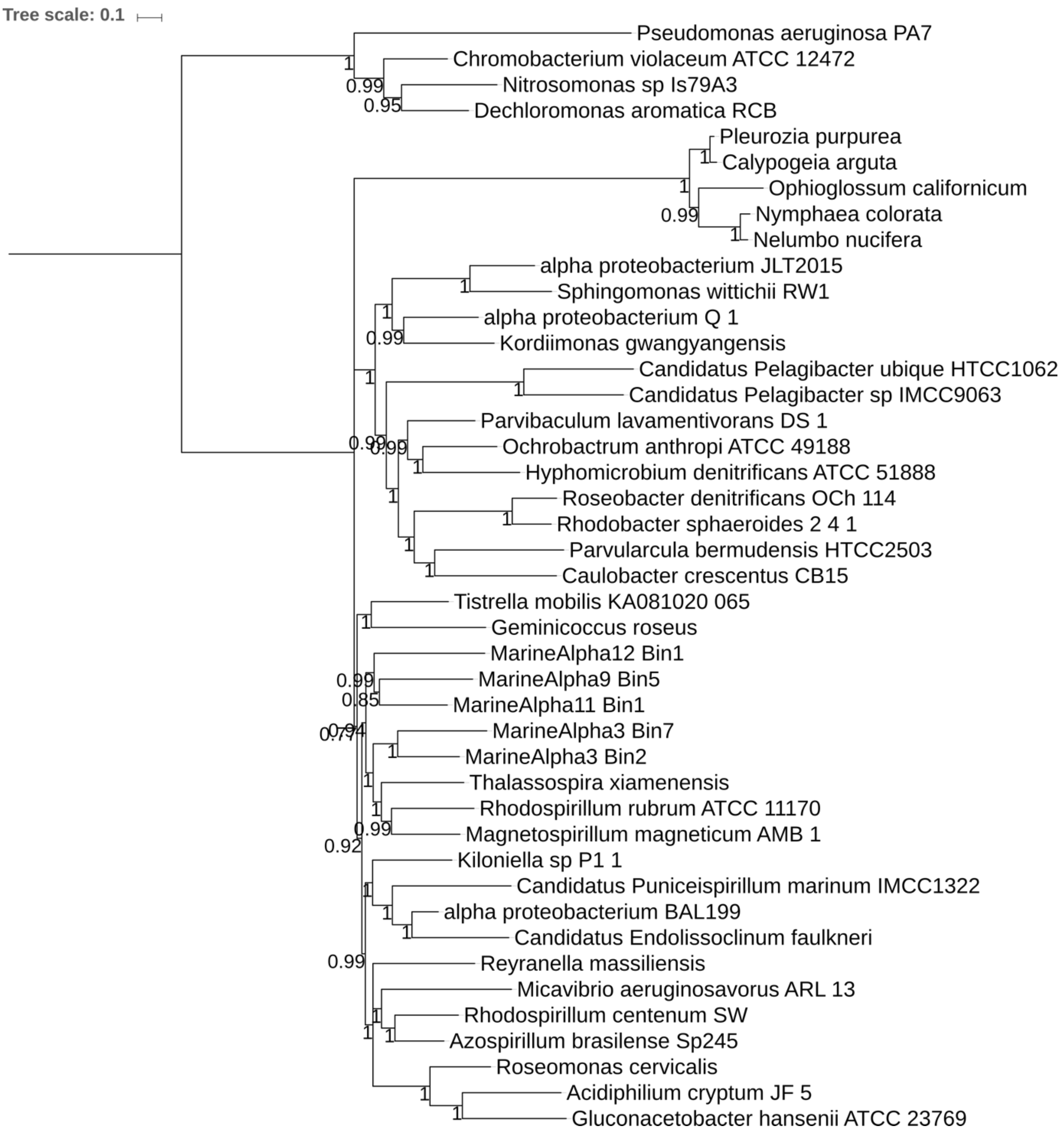


**Supplementary Fig. 18 | Bayesian phylogenetic tree of backbone alphaproteobacteria, Pelagibacterales and mitochondria in the 18-alphamitoCOGs dataset.** The tree is rooted with representatives of Beta- and Gammaproteobacteria. Node values show posterior probability support values.

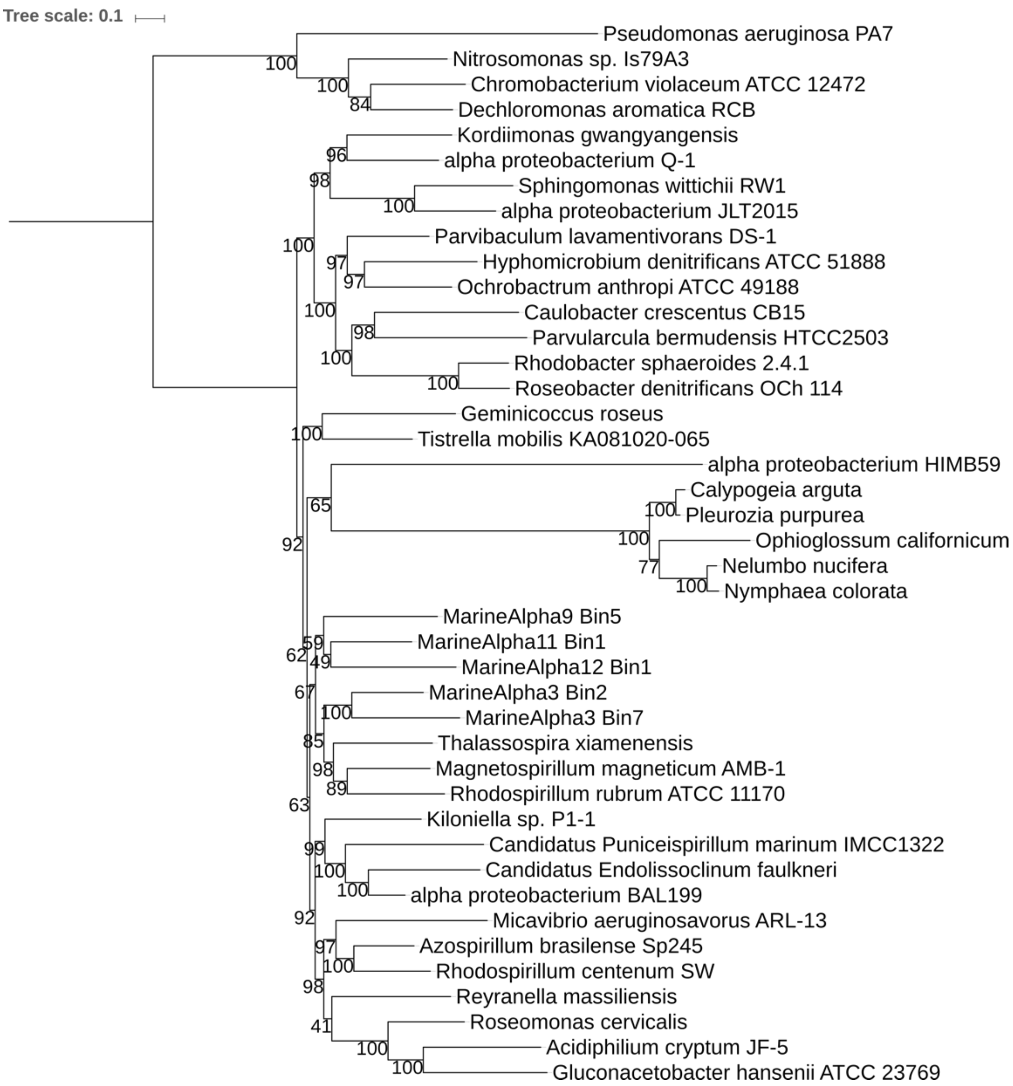


**Supplementary Fig. 19 | ML phylogenetic tree of backbone alphaproteobacteria, alphaproteobacterium HIMB59 and mitochondria in the 18-alphamitoCOGs dataset.** The tree is rooted with representatives of Beta- and Gammaproteobacteria. Node support values are based on the bootstrap results after 1000 iterations.

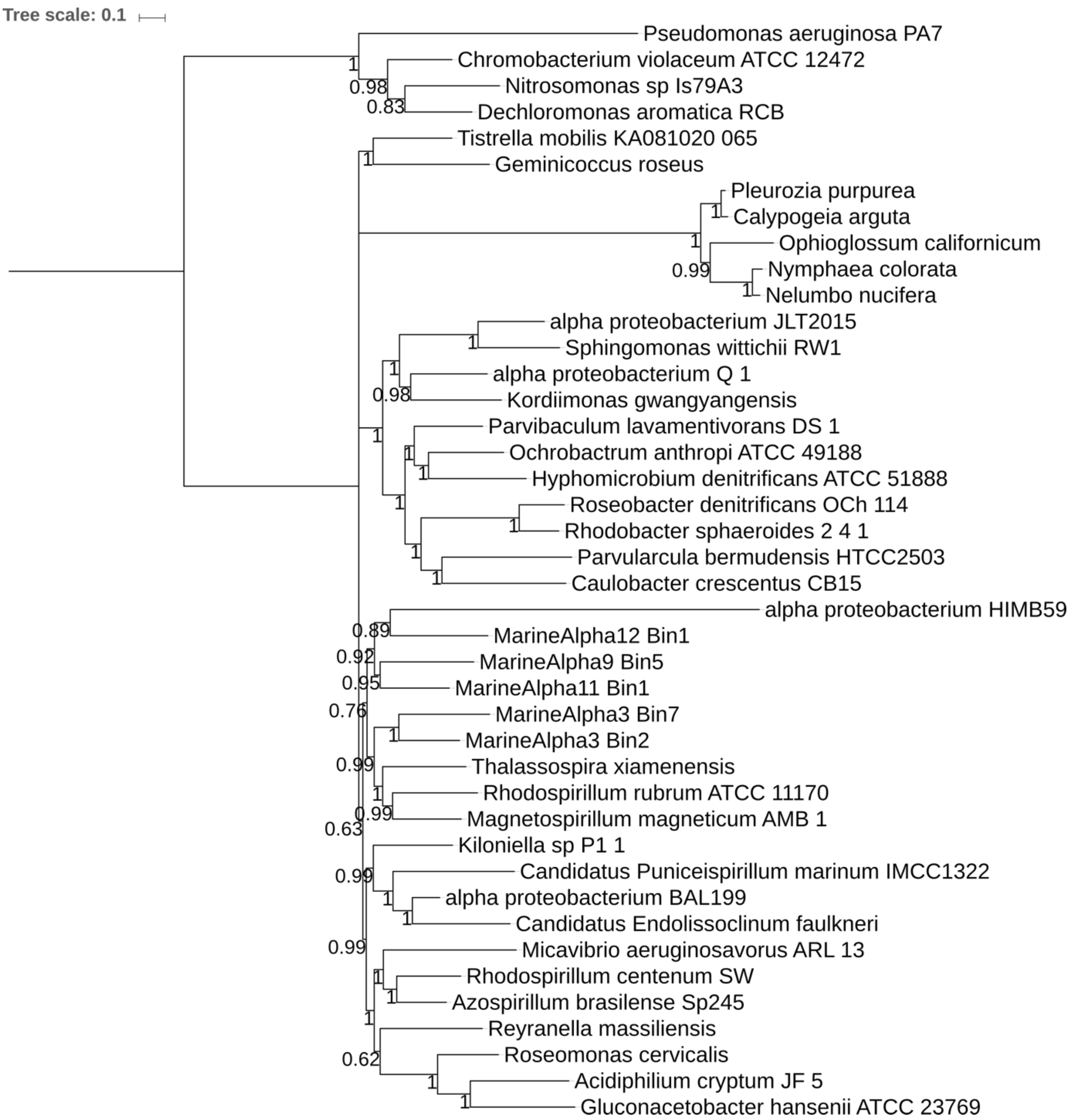


**Supplementary Fig. 20 | Bayesian phylogenetic tree of backbone alphaproteobacteria, alphaproteobacterium HIMB59 and mitochondria in the 18-alphamitoCOGs dataset.** The tree is rooted with representatives of Beta- and Gammaproteobacteria. Node values show posterior probability support values.

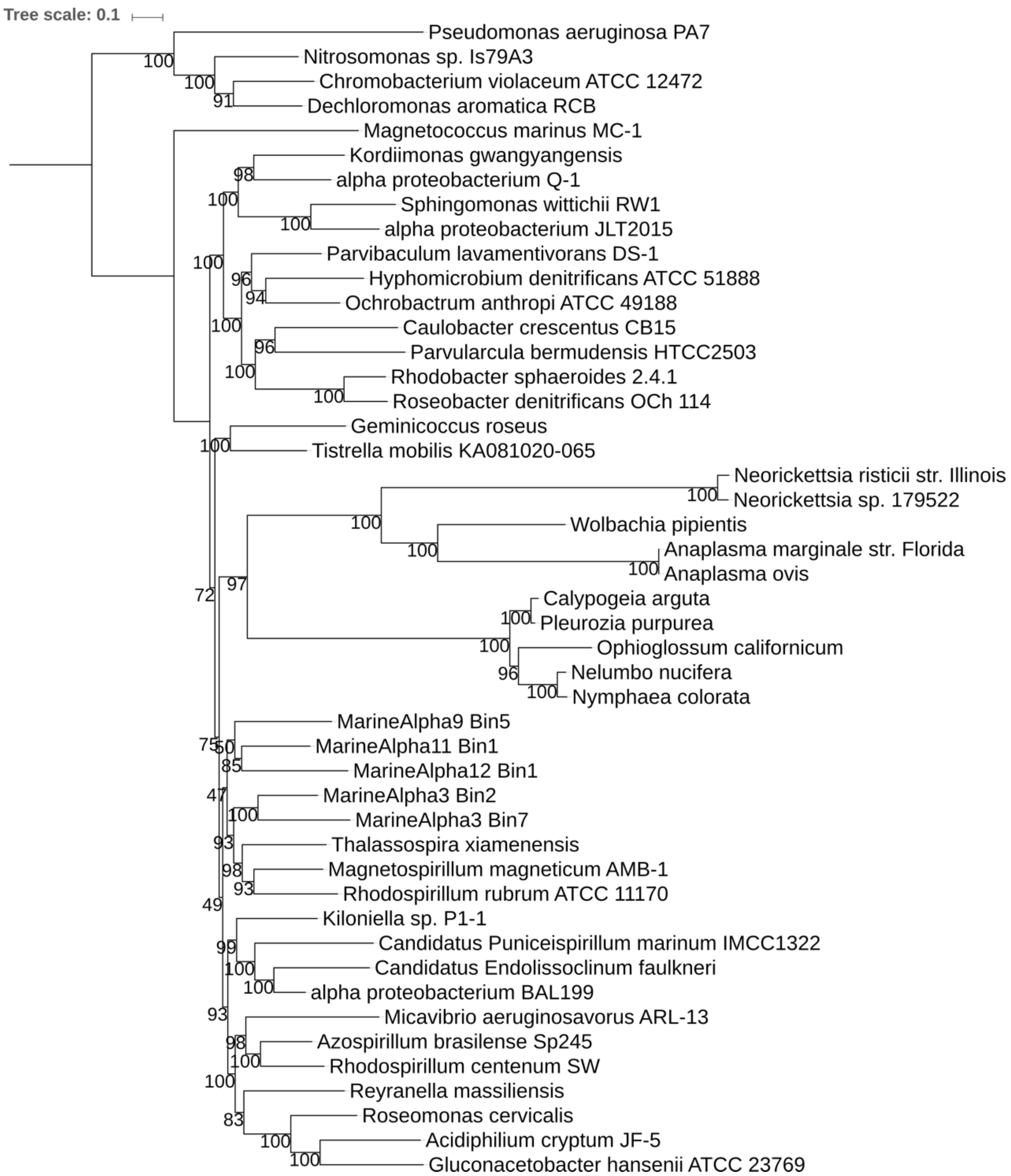


**Supplementary Fig. 21 | ML phylogenetic tree of backbone alphaproteobacteria, Rickettsiales and mitochondria in the 18-alphamitoCOGs dataset.** The tree is rooted with representatives of Beta- and Gammaproteobacteria. Node support values are based on the bootstrap results after 1000 iterations.

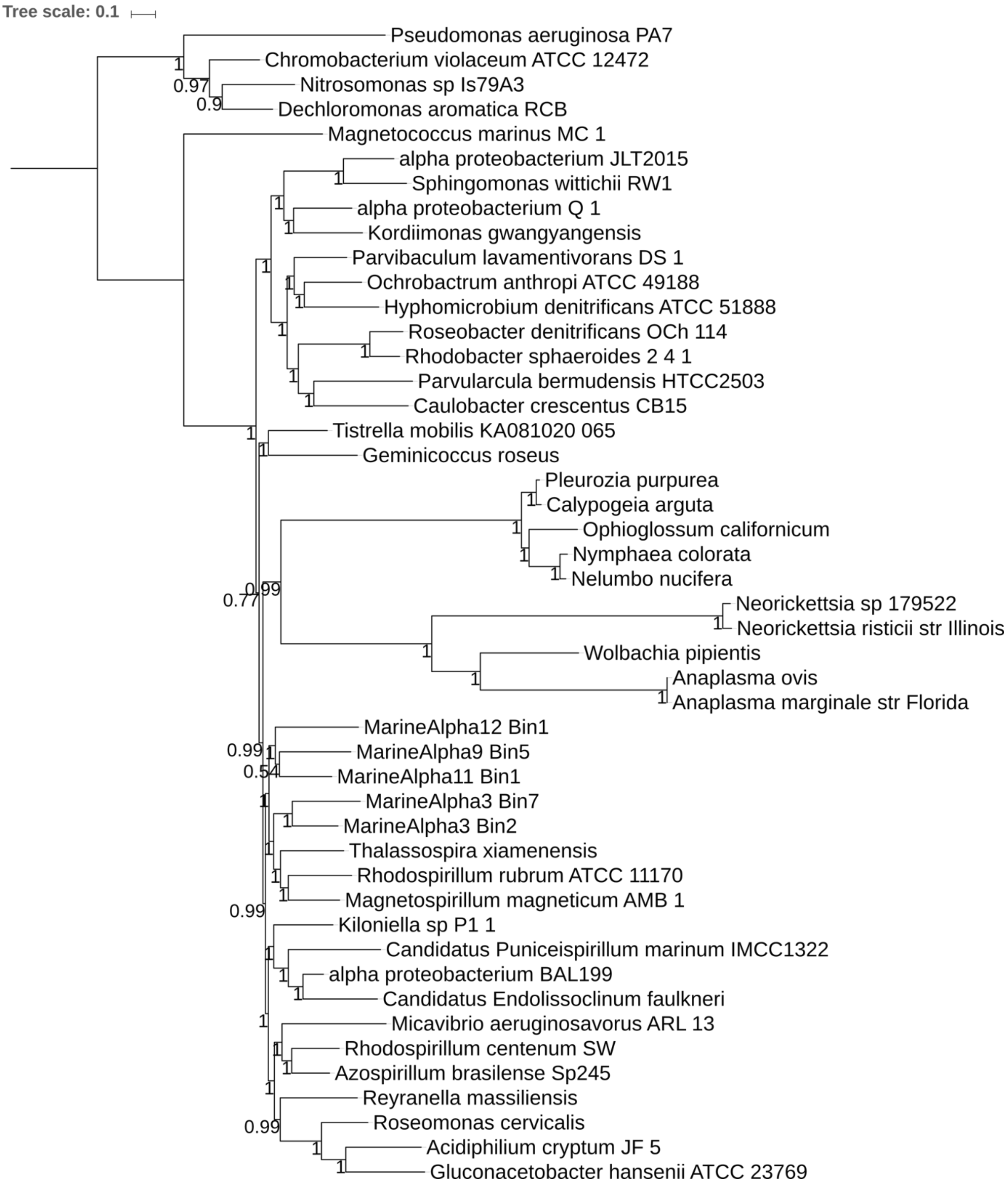


**Supplementary Fig. 22 | Bayesian phylogenetic tree of backbone alphaproteobacteria, Rickettsiales and mitochondria in the 18-alphamitoCOGs dataset.** The tree is rooted with representatives of Beta- and Gammaproteobacteria. Node values show posterior probability support values.

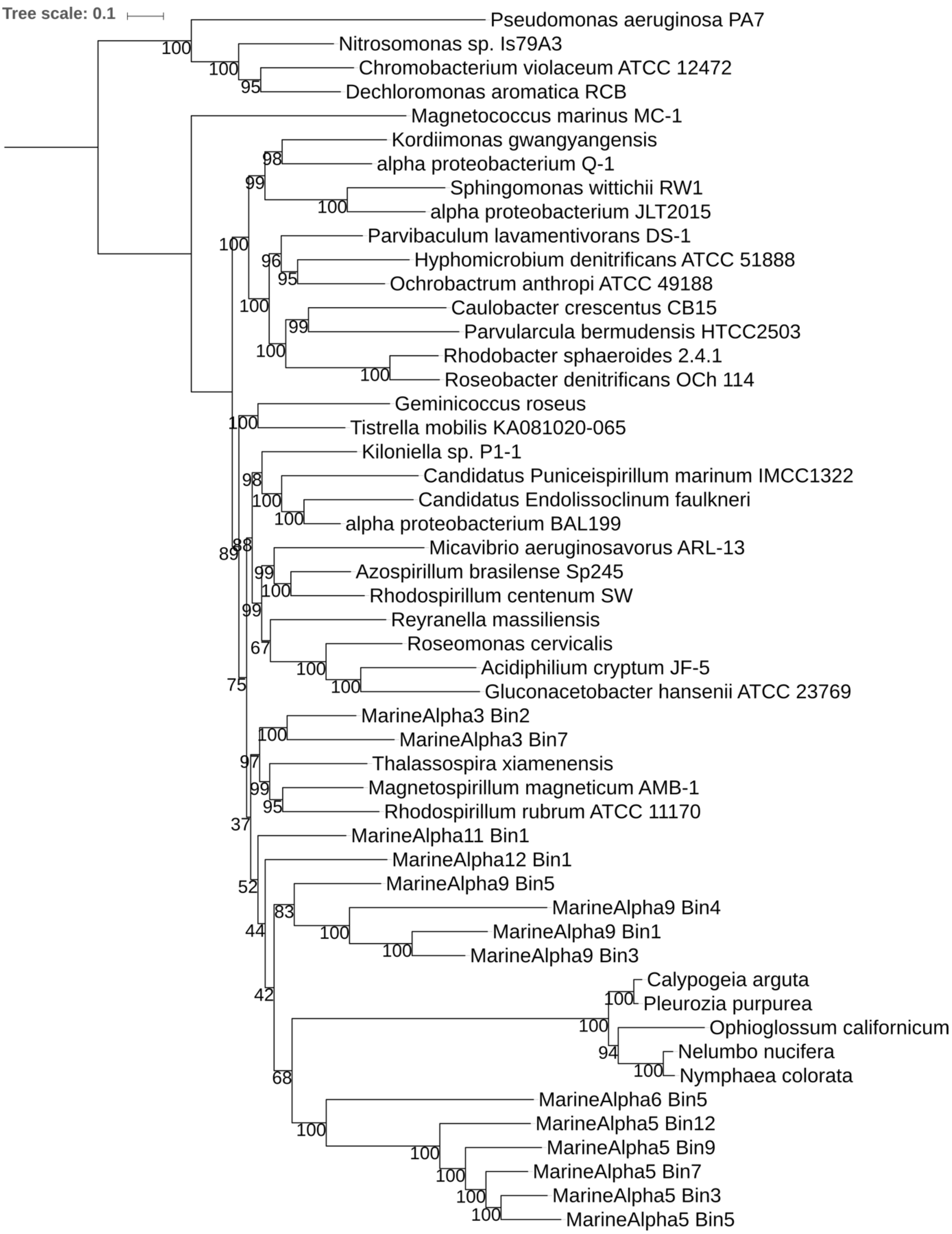


**Supplementary Fig. 23 | ML phylogenetic tree of backbone alphaproteobacteria, FEMAG I, FEMAG II and mitochondria in the 18-alphamitoCOGs dataset.** The tree is rooted with representatives of Beta- and Gammaproteobacteria. Node support values are based on the bootstrap results after 1000 iterations.

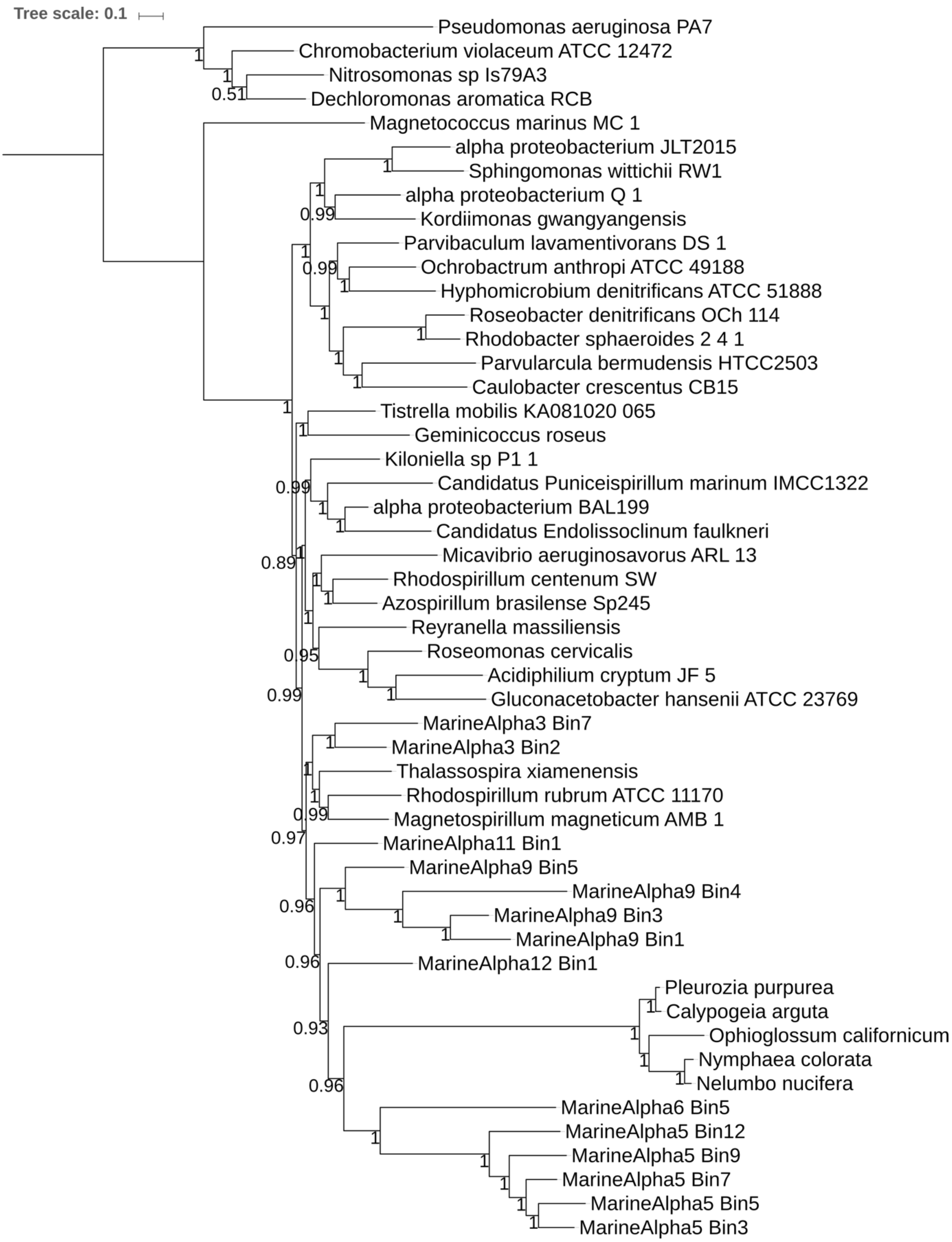


**Supplementary Fig. 24 | Bayesian phylogenetic tree of backbone alphaproteobacteria, FEMAG I, FEMAG II and mitochondria in the 18-alphamitoCOGs dataset.** The tree is rooted with representatives of Beta- and Gammaproteobacteria. Node values show posterior probability support values.
